# The *Drosophila* EcR-Hippo component Taiman promotes epithelial cell fitness by control of the Dally-like glypican and Wg gradient

**DOI:** 10.1101/2024.03.31.587486

**Authors:** Colby K. Schweibenz, Victoria C. Placentra, Kenneth H. Moberg

## Abstract

Rapidly dividing cells can eliminate slow growing neighbors through the apoptotic process of cell competition. This process ensures that only high fitness cells populate embryonic tissues and is proposed to underlie the ability of oncogene-transformed cells to progressively replace normal cells within a tissue. Patches of cells in the *Drosophila* wing disc overexpressing the oncogenic Taiman (Tai) transcriptional coactivator kill normal neighbors by secreting Spz ligands that trigger pro-apoptotic Toll signaling in receiving cells. However, extracellular signaling mechanisms responsible for elimination of slow growing cells by normal neighbors remain poorly defined. Here we show that slow growing cells with reduced Tai (Tai^low^) are killed by normal neighbors through a mechanism involving competition for the Wingless (Wg/Wnt) ligand. Elevated Wg signaling significantly rescues elimination of Tai^low^ cells in multiple organs, suggesting that Tai may normally promote Wg activity. Examining distribution of Wg components reveals that Tai promotes extracellular spread of the Wg ligand from source cells across the wing disc, thus ensuring patterned expression of multiple Wg-regulated target genes. Tai controls Wg spread indirectly through the extracellular glypican Dally-like protein (Dlp), which binds Wg and promotes its extracellular diffusion and capture by receptors. Data indicate that Tai likely controls Dlp at two levels: transcription of *dlp* mRNA and Dlp intracellular trafficking. Overall, these data indicate that the Tai acts through Dlp to enable Wg transport and signaling, and that cell competition in the Tai^low^ model arises due to inequity in the ability of epithelial cells to sequester limiting amounts of the Wg growth factor.

## Introduction

The development of an organism is a tightly controlled process, with multiple signaling pathways converging to achieve correct patterning, morphogenesis, and cell fates. As the earliest cells begin to populate tissues, it is advantageous that the fittest progenitors give rise to the entire organism. One example of this is in the formation of the epiblast—early embryonic cells go through a selection process, where more fit cells with higher amounts of pluripotency factors such as TEAD or Myc outcompete cells with a relatively lower dose of these factors[1,2]. This process of pruning unfit cells and selection of fitter cells for survival is known as “cell competition.”

The phenomenon of cell competition was first described in the context of *Drosophila melanogaster* cells with defective ribosomal protein function, termed “*Minutes*”, that could survive when surrounded by like cells but died in a heterotypic environment mixed with wildtype cells[3]. Cells that die in a heterotypic environment are called “losers,” as they undergo apoptosis through mechanisms triggered by neighboring cells termed “winners”. A variety of apoptotic mechanism have been proposed to underlie loser fate, including exposure to the pro-apoptotic cytokine Spätzle (Spz) produced by oncogene transformed ‘winners’[4–6], non-autonomous induction of autophagy[7,8], loss of apicobasal polarity[9], and inequities in expression of the transmembrane factor Flower[10], the secreted protein SPARC[11], or the Ca^+2^ binding protein Azot[12]. Altering the dose of the dMyc oncogene has also been proposed to generate competitive differences by altering the ability of cells to compete for limiting amounts of the Dpp morphogen[13,14].

In previous work we found evidence that wing cells overexpressing the conserved coactivator and oncogene Taiman (Tai; human AIB/SRC3/NCOA3) kill neighboring cells by overproduction and secretion of Spz ligands, in particular Spz4[6], which is similar to a proposed mechanism by which excess Myc confers winner status[4]. Intriguingly, *Drosophila* cells with reduced Tai expression are viable, proliferate more slowly than normal cells, and generate small but properly patterned organs [15]. However, they have not been tested for their fitness status, and whether they become losers that resemble cells with reduced levels of the dMyc oncogene (i.e., Myc^low^ cells, as in [4,5]). Here we have used a homozygous viable allele of *tai* that behaves as a weak hypomorph to investigate the role of Tai in cell competition. We find that larval wing disc cells with reduced Tai (Tai^low^) survive in a homotypic environment but are eliminated by neighboring wildtype cells in a heterotypic environment of clonal mosaics. A genetic screen to identify pathways involved in competitive elimination of Tai^low^ cells identifies alleles of the Wnt/Wg pathway inhibitor *Adenomatous polyposis coli* (*APC*) as suppressors of Tai^low^ elimination, suggesting that Tai^low^ elimination may be due to a deficit of Wg signaling. Examining the Tai-Wg link more closely reveals that Tai is required for formation of the Wg gradient in the larval wing pouch and for patterned activation and repression of Wg target genes at different points along the Wg gradient. We trace this defect to a role for Tai in promoting levels and intracellular trafficking of the Dally-like protein (Dlp) glypican, which binds extracellular Wg and facilities its movement away from source cells. Evidence suggests that Tai is a key determinant of the effect of Dlp on Wg steady-state levels: excess Dlp triggers Wg accumulation in Tai-depleted cells, but triggers Wg loss in cells that also express Tai. Dlp accumulates intracellularly in Tai^low^ clones, confirming a link between Tai and Dlp trafficking. Given that cells lacking *Apc* are ‘winners’ in the wing pouch and adult intestine[16,17], we propose that Tai regulates competitive status though an underlying developmental role in promoting Dlp-dependent Wg capture and signaling. As a result, cells with reduced Tai are disadvantage with normal neighbors for capture of limiting amounts of Wg.

## RESULTS

### Cells with reduced Tai are selectively killed in clonal mosaics

Combining the weak, viable hypomorph *tai^k15101^* with a deletion (*Df ED678*) that completely removes the second copy of *tai* produces small but normally patterned adults with proportionally smaller organs[15]. Larval wing discs from these *taik15101/Df* animals contain background levels of apoptotic cells (**Fig. 1A-A’,D**) suggesting that wing size reduction is the result of a proliferative deficit rather than surplus apoptosis. This hypothesis is consistent with our previous finding that Tai promotes cell division in the larval wing[15]. Generating boundaries between *tai^k15101^* cells (*tai^k15101^,FRT40A*) and normal cells (*FRT40A*) using the Flp-FRT mosaic system results in significantly elevated cleaved Dcp-1 (cDCP-1) caspase within *tai^k15101^* clones (**Fig. 1B-B”, C-C”, D**), indicative of apoptosis. This excess cDcp1 is especially evident in *tai^k15101^*clones located in the wing pouch, a developing epithelium that is widely used to assess competitive status [4,18]. These data show that pouch cells with one hypomorphic *tai* allele (i.e., *tai^k15101^/Df*) are viable when surrounded by like cells, but that cells with two copies of the *tai^k15101^* hypomorph die when surrounded by *tai^wt^*cells. These data indicate the existence of an extracellular competition mechanism that allows normal *tai^wt^* cells to kill *tai^k15101^*(hereafter *tai^low^*) neighbors.

**Figure 1.**
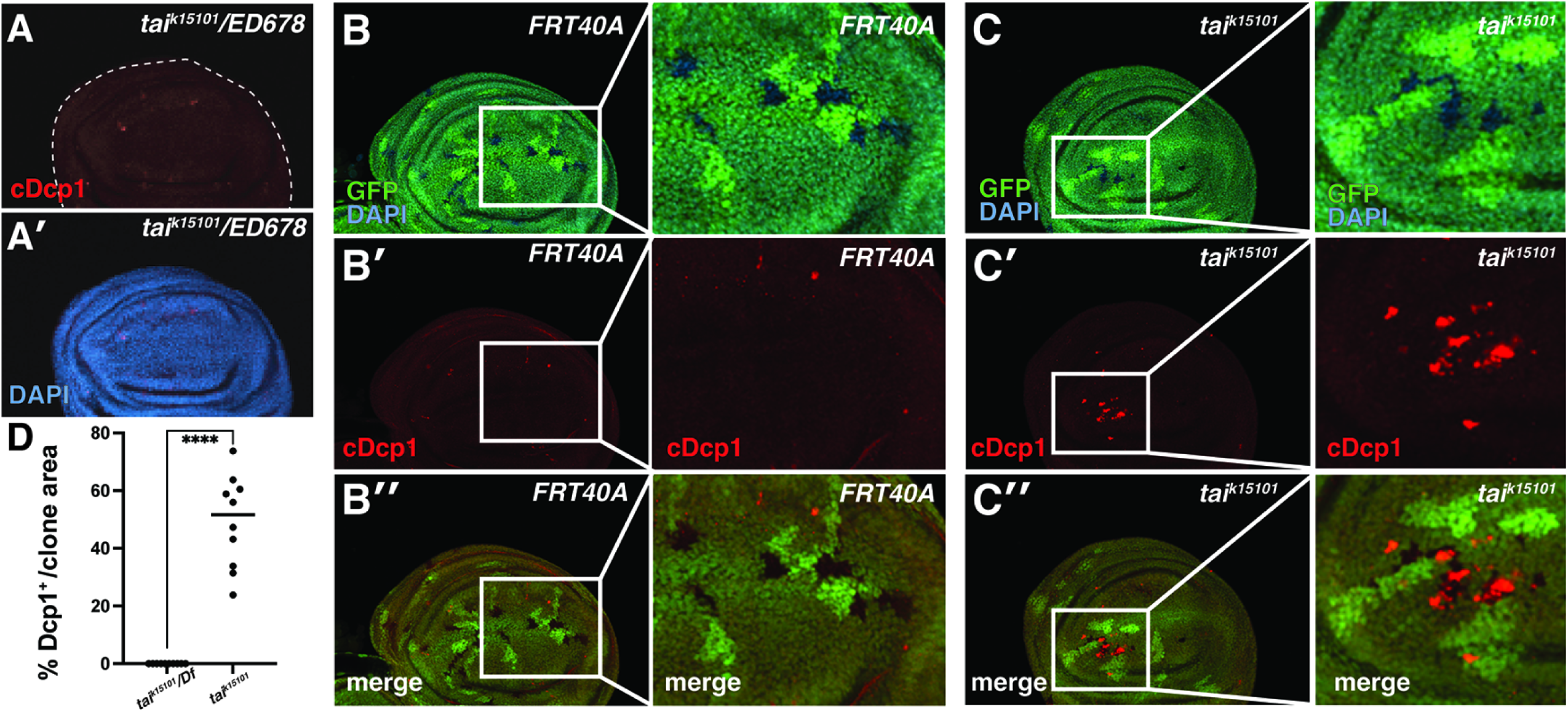
*tai^low^*cells survive in a homotypic environment but die in a heterotypic environment. **(A-A’)** 3^rd^ larval instar *tai^k15101^/Df(2L)ED678* wing disc stained with anti-cleaved Dcp-1 (cDcp1; red) and nuclei (DAPI; blue). (**B-C**) *hsFlp* generated clones of control *FRT40A* (**B-B”**) or *tai^k15101^,FRT40A* (**C-C”**) cells marked by the absence of GFP (green) and co-stained for cDcp-1 (red) and nuclei (DAPI; blue). Magnified insets are provided. **(D)** Quantification of cDcp-1 in *tai^k15101^/Df(2L)ED678* heterotypic discs or *tai^k15101^* clones calculated by percent Dcp1-positive area/total clone area (Student t-test, *p* <0.0001).

### Elimination of *tai^low^* cells requires Wg, Hippo, and canonical apoptosis pathways

A *mini-w^+^*minigene present in the *tai^k15101^* hypomorph generates red pigment that visually marks *tai^low^* heterozygous or homozygous cells in the adult eye. As in the larval wing disc, *tai^k15101^/Df* cells readily populate an adult eye composed of alike cells, but patches of *tai^low^* cells form small clones when placed in competition with control (*white^-^*;*FRT40A*) cells using an eye-specific Flpase transgene (*eyFLP*) (**Fig. 2A-B**). By comparison, cells carrying a *mini-w^+^* marked *FRT40* chromosome occupy approximately 50% of the adult eye.

**Figure 2.**
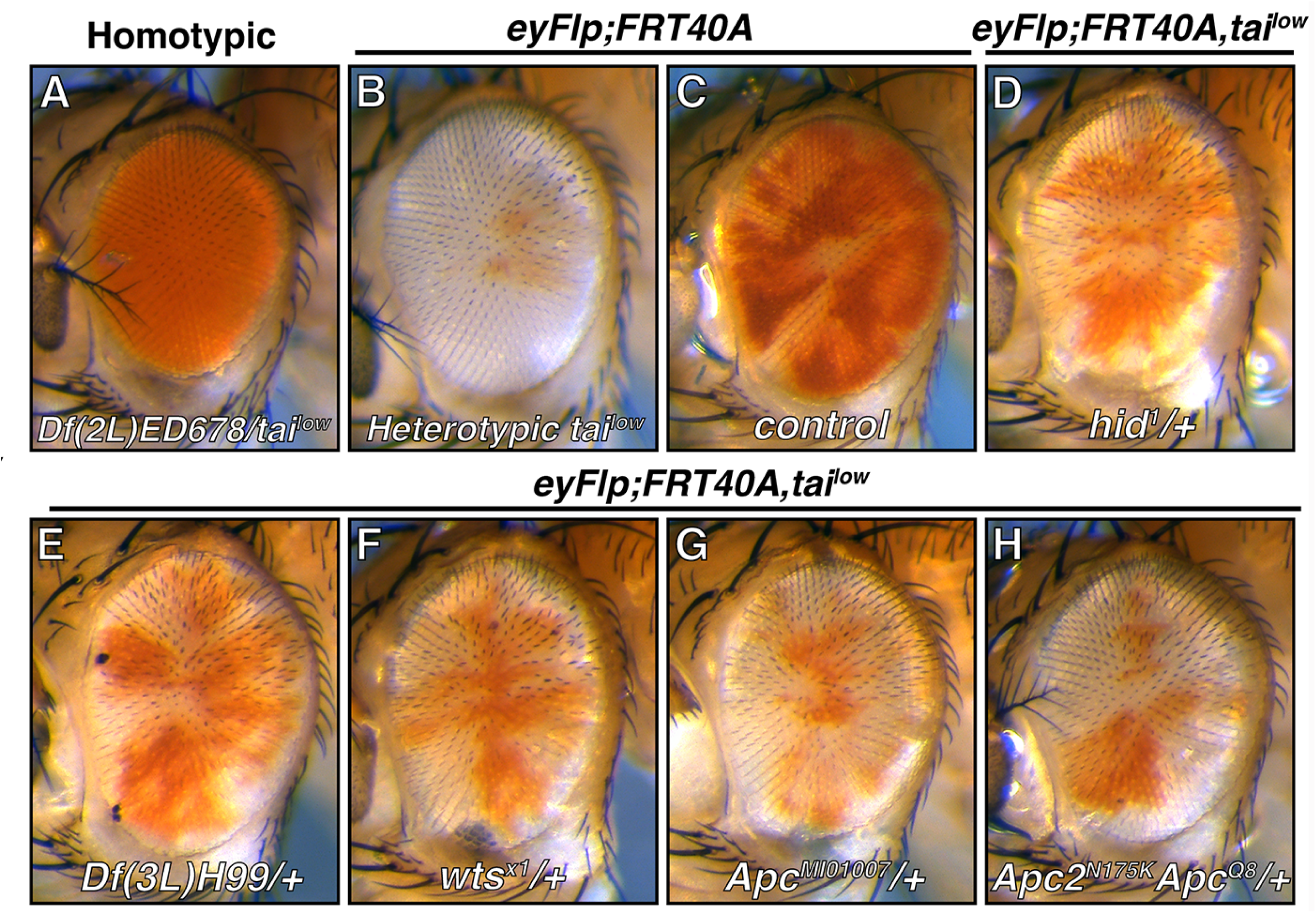
Modification of *tai^low^*clone survival in the adult eye. Adult female eyes of the indicated genotypes: (**A**) *tai^k15101^/Df(2L)ED678*; (**B-C**) *eyFLP* mosaic eyes with *w-;FRT40A* flipped over (**B**) *tai^k15101^,FRT40A* or (**C**) *FRT40A,m-w^+^*; (**D-H**) *eyFLP* mosaic eyes with *tai^k15101^;FRT40A* flipped over *FRT40A,m-w^+^*in the background of (**D**) *hid^1^*, (**E**) *Df(3L)H99*, (**F**) *wts^x1^*, (**G**) *Apc^MI01007^*, or (**H**) *Apc2^N175K^,Apc^Q8^*.

To identify genes and pathways involved in elimination of *tai^low^*eye cells, a collection of mutant alleles of candidate competition factors (**Supplemental Table 1**) were screened for dominant rescue of *tai^low^*small clone size in the adult eye (**Fig. 2C-H**). This approach yielded ‘hits’ in the pro-apoptotic Rpr-Hid-Grim (RHG) factors (*hid^1^* and the *H99* deletion), the Hippo pathway kinase *warts* (*wts^1^*), and the Wg pathway inhibitor *Adenomatous polyposis coli* (*Apc^MI01007^*). Alleles affecting components of the Spz-Toll, IMD, or ecdysone pathways did not obviously modify *tai^low^* clone size, nor did alleles of other candidate factors such as the transmembrane proteins *flower* and *Sas*, the transcription factors *p53* and *Xrp1,* the autophagy factor *Atg1*, and the secreted glycoprotein *Sparc* [8-11,19-22] (data not shown). These genetic data confirm separate elements of *tai^low^*loser fate: rescue by *hid^1^* and *H99* provide genetic evidence that elimination of *tai^low^* cells requires the canonical RHG-Diap-Caspase pathway (rev. in [23]), while rescue by *wts^1^* is consistent with Tai protecting against the “loser” fate through Yorkie, which is bound by Tai and phosphorylated by Wts[15,24,25]. By contrast, *tai^low^*rescue by *Apc* suggests an unexpected link between Tai competitive status and Wg signaling. A second *Apc* mutation (*Apc^Q8^*) in combination with an inactivating allele in its paralog *Apc2* (*Apc2^N175A^*) similarly increases survival of *tai^low^* cells in the eye (**Fig. 2H**).

The larval wing pouch is bisected along the D-V (dorsoventral) midline by a stripe of Wg-expressing cells that generate mirror image gradients (dorsal and ventral) that guide wing growth and patterning[26]. Quantification of *tai^low^* clone size within the larval wing pouch at 50hrs post induction (*hsFlp*) indicates that *RHG*, *wts*, or *Apc/Apc2* alleles recovered in the eye screen also increase the size of *tai^low^* pouch cells (**Figure 3A-H**), with *H99* and the compound *Apc^Q8^,Apc2^N175A^* allele having the largest effects. *H99* and *Apc* alleles also enhance recovery of 76hr old *tai^low^* clones, which are normally eliminated by this time point (**Fig. S1**). Together, these data indicate that competitive elimination of *tai^low^* cells requires RHG proteins and can be attenuated by derepressing the Wg or Hippo pathways.

**Figure 3.**
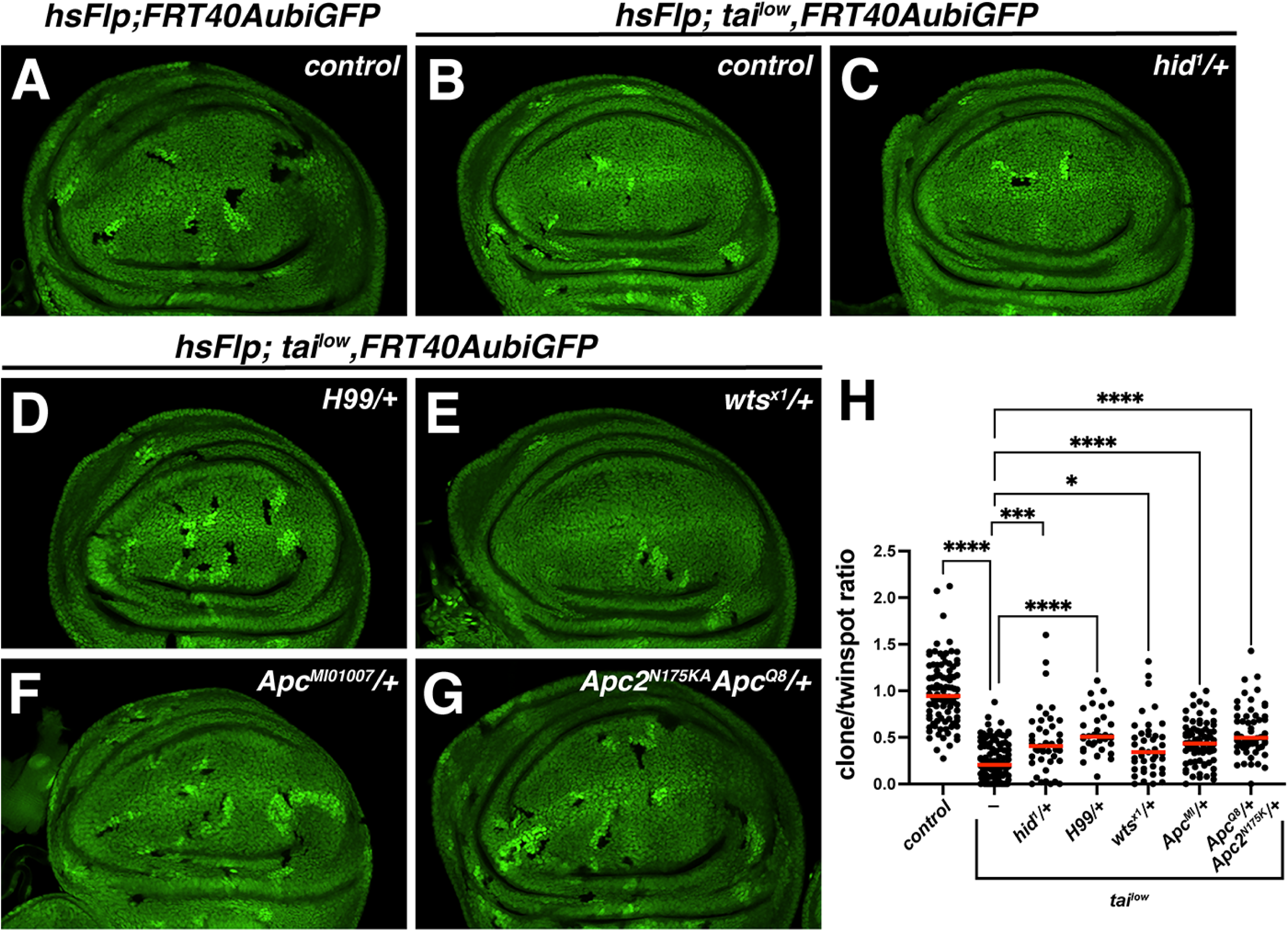
Alleles from the eye screen also modify *tai^low^* clone survival in L3 wing pouch. (**A-G**) Larval wing discs bearing *hsFlp/FRT*-generated 50hr-old *tai^low^* clones (GFP-negative) either (**B**) alone, or in the background of the indicated alleles (**C-G**). Twinspots appear brighter due to two copies of *GFP*. **(H)** Quantification of *control*(*FRT40A*) or *tai^low^* size ratio (area ratio of clone:twin-spot) alone (-) or in the indicated genetic backgrounds.. \*\*\*\**p* <0.001, \*\*\**p*= 0.0003, \**p* = 0.0157, *ns* = not significant (One-way ANOVA with Dunnett post hoc test).

### Tai modulates transcription of Wg-regulated genes and formation of the Wg gradient

The extended survival of 76hr *tai^low^* clones in *Apc* and *Apc,Apc2* heterozygous backgrounds imply a relatively strong link between Tai and Wg signaling. To test sensitivity of Wg pathway activity to Tai dosage, *UAS* transgenes were used to test whether *tai* overexpression or reduction affects transcription of two Wg target genes: *naked cuticle* (*nkd*), a mid-threshold target of the Wg gradient that is expressed in bands of cells on either side of the DV midline[27] (*nkd-lacZ*), and *wg* itself, which is repressed by an autoinhibition loop that refines *wg* transcription to a narrow stripe at the DV midline[28] (*wg-lacZ*). Tai overexpression (*en>tai*) expands *nkd-lacZ* expression into dorsal and ventral domains that correspond to distal tails of the Wg gradient and normally express low *nkd* (**Fig. 4A-A“, B-B”**, see brackets, and **4D-E**). Tai depletion strongly retards *nkd-lacZ* in regions close to the DV midline that normally receive high Wg (**Fig. 4C-C”, D-E**). Examining *wg* transcription with the *wg-lacZ* transgene shows that *tai* RNAi (*enGal4*) causes *wg-lacZ* to spread dorsally and ventrally away the DV margin (**Fig. 5B-B’**, see arrow), while reciprocally elevating *tai* has little effect on *wg* transcription (**Fig. 5C-C’**). These data are consistent with Tai acting as a positive upstream regulator of Wg signaling in pouch cells, such that activation of *nkd* and autoinhibition of *wg* each fail when Tai is removed.

**Figure 4.**
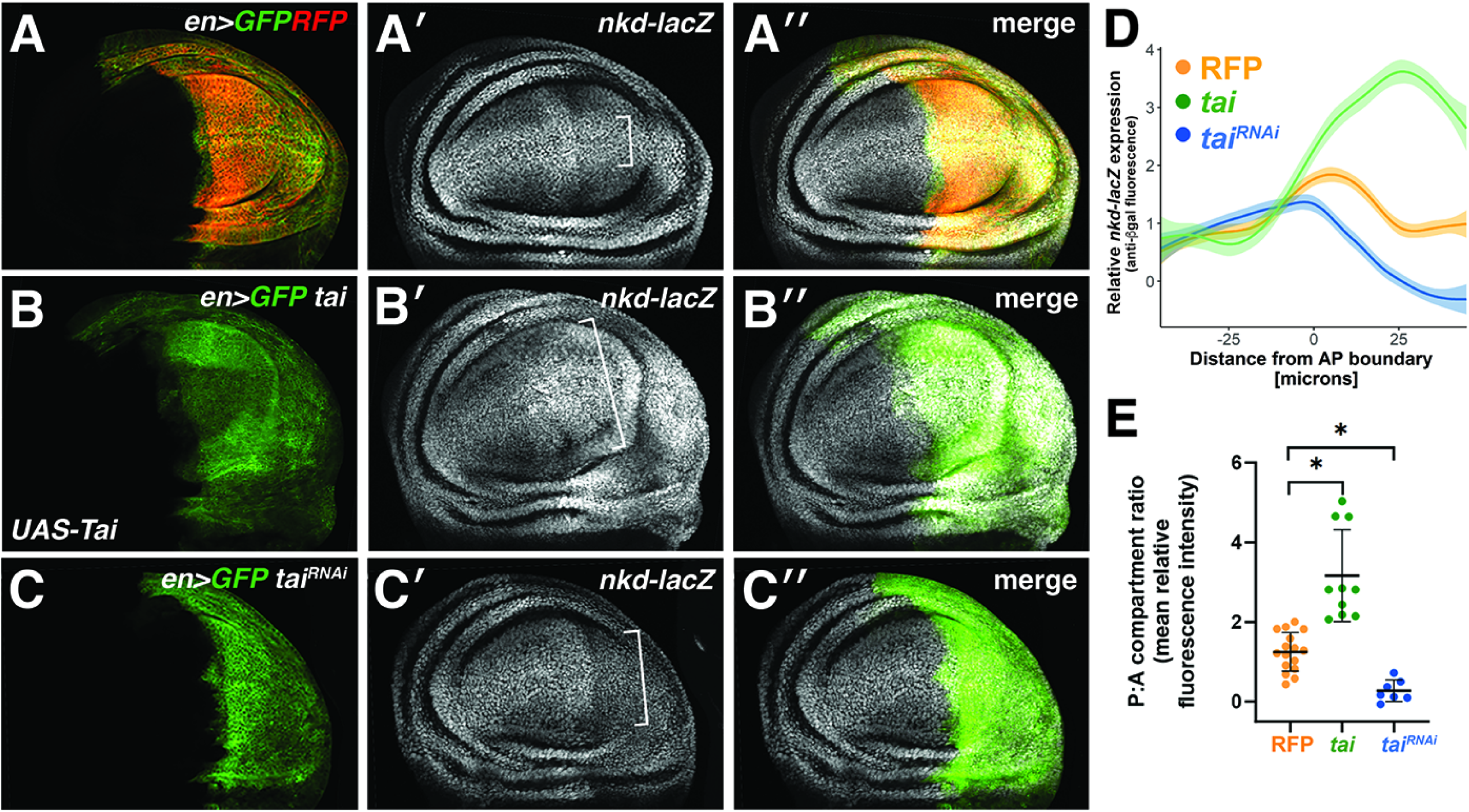
*tai* promotes expression of the Wg pathway reporter *nkd-lacZ*. (**A-C**) Expression of the *nkd-lacZ* reporter (greyscale) in (**A-A”**) *control* (*en>GFP,RFP*), (**B-B”**) Tai overexpressing (*en>tai,GFP*), or (**C-C”**) Tai-depleted larval wing discs. **(D)** Relative anti-ßGal fluorescence intensity on either side of the A-P boundary among pouch cells located just below the D/V boundary. **(E)** Mean relative fluorescence intensity plotted as a ratio of posterior to anterior intensity. *RFP* vs *tai* *p=0.0004; *RFP* vs *tai*^RNAi^ \**p*<0.0001 (Unpaired student t-test with Welch’s correction).

**Figure 5.**
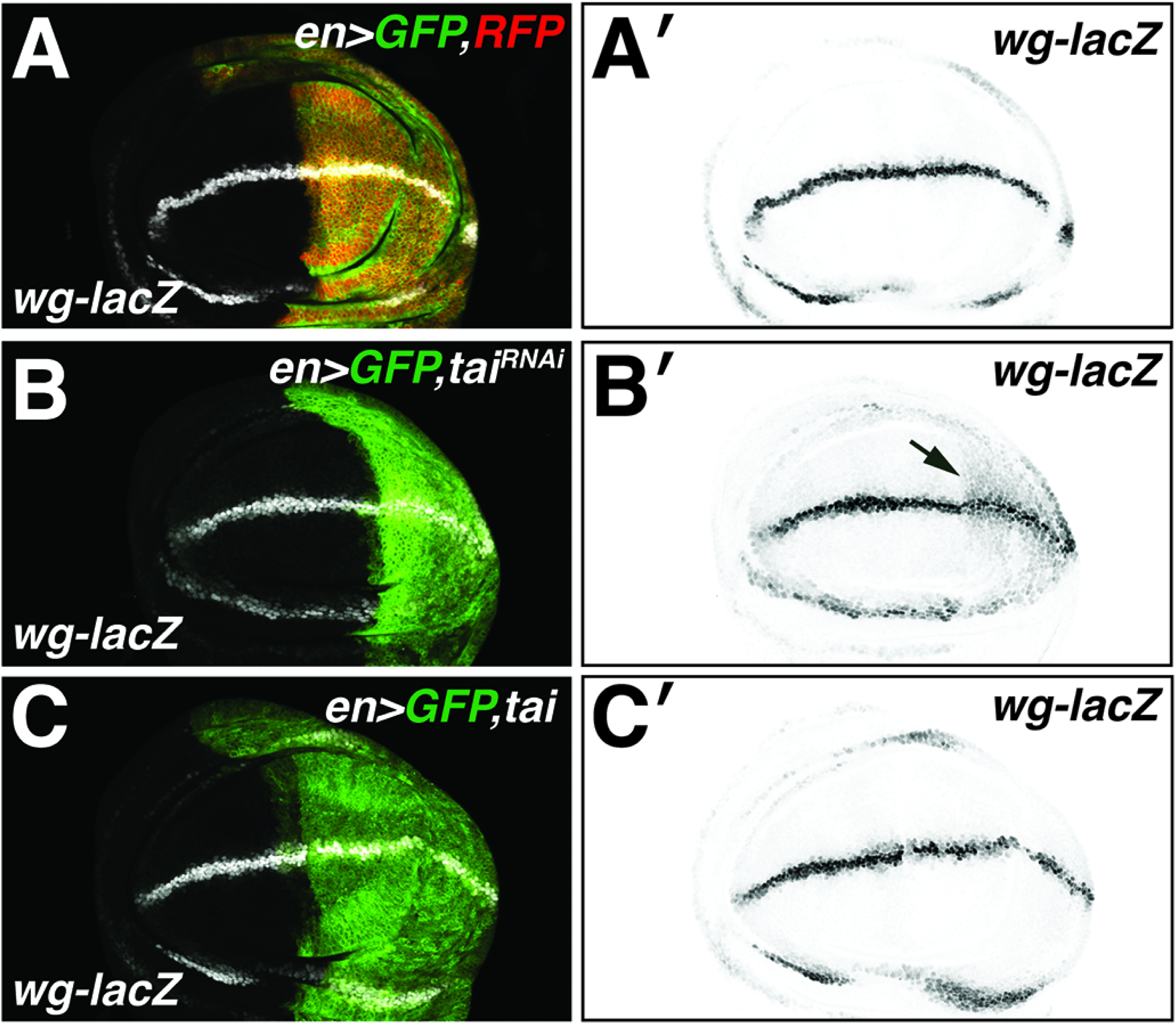
Tai is required for the auto-inhibitory loop that refines *wg* transcription. **(A-C)** Expression of the *wg-lacZ* reporter in *enGal4* larval wing discs as detected by anti-ßGal staining (white in **A-C**, black in **A’-C’**) in *control* RFP (**A**), Tai-depleted (**B**), or Tai overexpressing (**C**) discs. Arrow in **B’** indicates failure to refine *wg-lacZ* to the DV boundary in P-domains depleted of Tai.

Failure to constrain *wg-lacZ* activity to DV margin cells in Tai-depleted discs (see **Fig. 5B’**) resembles the effect of mutations that block receipt of the Wg ligand[28]. Examination of the Wg gradient in pouch cells with reduced Tai in the posterior domain (*en>tai* RNAi) detects robust Wg in source cells but significant reduction in Wg spreading into the dorsal and ventral pouch (**Fig. 6A** vs. **6C**, arrows). This loss of the Wg gradient occurs despite expansion of the *wg* expression domain in Tai RNAi pouches (see **Fig. 5B-B’**). Overexpression of Tai produces gaps in the stripe of Wg protein in DV cells (**Fig. 6B**, arrows) that also does correlate with a loss of *wg* transcription (see **Figs. 5C-C’**), suggestive of a post-transcriptional effect of excess Tai on Wg stability, uptake, or trafficking in the DV regions. Quantification of anti-Wg staining intensity confirms that Tai RNAi blocks formation of the Wg gradient and indicates that Tai overexpression elevates Wg in regions further from the DV margin (**Figs. 6D-E**). Considered together, these data are consistent with a requirement for Tai in forming the Wg gradient by control of Wg protein extracellular diffusion and/or internalization and turnover.

**Figure 6:**
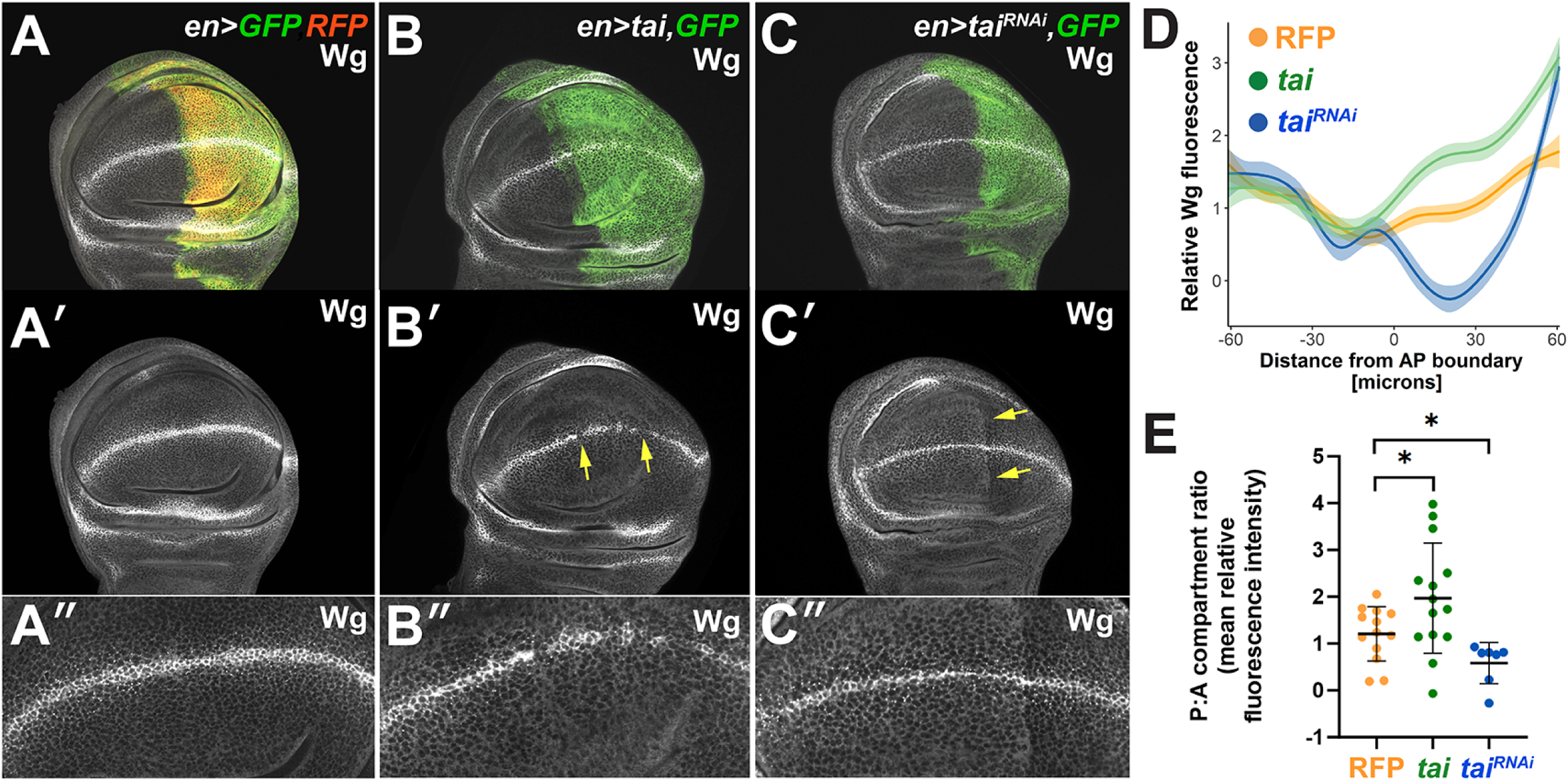
Tai is required for formation of the Wg gradient. Wg protein (greyscale) in (**A-A”**) *control* (*en>GFP,RFP*), (**B-B”**) Tai overexpressing, or (**C-C”**) Tai-depleted larval wing discs. *enGal4* activity in the P-domain is marked by GFP. Arrows in **B’** mark gaps in Wg DV stripe characteristic of Tai overexpression. Arrows in **C’** mark the clear drop in total Wg on the DV flanks of P-domains depleted of Tai. **A”-C”** provide higher magnification views of regions in **B’-C’. (D)** Relative Wg fluorescence intensity of pouch cells below D/V boundary (to avoid Wg source cells) plotted from anterior to posterior. **(E)** Mean Wg relative fluorescence intensity plotted as a ratio of posterior to anterior intensity. *RFP* vs *tai* \**p*=0.0443, *RFP* vs *tai^RNAi^* \**p*<0.0156 (Unpaired student t-test with Welch’s correction).

### Tai is required for expression of Dally-like protein, a glypican required for Wg signaling

The GPI-anchored glypican Dally-like protein (Dlp) binds lipidated Wg and facilitates its extracellular diffusion and capture by Frizzled receptor[29–33]. Notably, homozygous loss of *Apc* produces competitive ‘winners’ by upregulating the secreted enzyme Notum, which then cleaves Dlp and deprives normal neighbors of the ability to sequester or sense Wg[17]. Given that *Apc* heterozygosity rescues *tai^low^*elimination, *tai* alleles were tested for modulation of Dlp levels in the larval pouch. Excess Tai significantly elevates Dlp protein among posterior cells (*en>tai*) and Tai depletion had the opposite effect of depleting Dlp (**Figs. 7A-C** and graphs in **7D-E)**. Expression of the Dlp-related glypican Dally[34] also rises upon Tai overexpression but is insensitive to Tai depletion (*dally-lacZ*; **Figs. S2A-C**), other than shrinkage of the *dally-lacZ* pattern due to reduced P-domain size due to Tai loss (arrows **Figs. S2A’** vs. **S2C’**). These data indicate that Tai is rate-limiting for Dlp expression in the pouch and may be able to induce *dally* expression when overexpressed.

**Figure 7:**
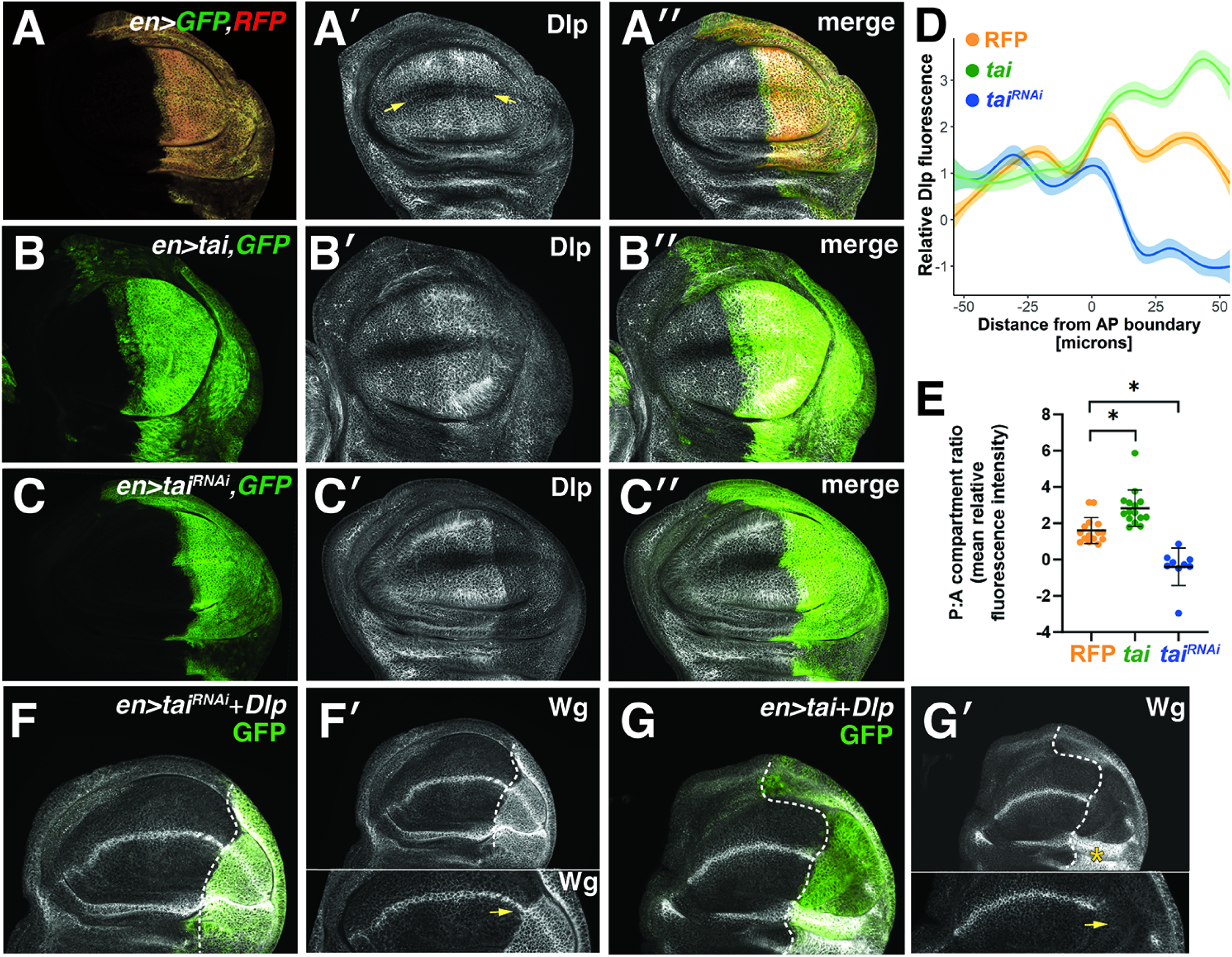
Tai controls Dally-like protein (Dlp) levels. **(A-C,F-G)** Anti-Dlp staining (greyscale) in (**A-A”**) *control* (*en>GFP,RFP*), (**B-B”**) Tai overexpressing (*en>tai,GFP*), and (**C-C”**) Tai-depleted cells. Arrows in **A’** mark Dlp exclusion from the DV boundary region. Overexpression of Dlp restores Wg in Tai-depleted cells (**F-F’**; see arrow in magnified view), but causes complete loss of the Wg DV stripe in Tai overexpressing disc regions (**G-G’**; see arrow in magnified view). **(D)** Relative Wg fluorescence intensities of pouch regions below D/V boundary plotted from anterior to posterior. **(E)** Mean relative Wg fluorescence intensity plotted as a ratio of posterior to anterior intensity. *RFP* vs *tai* \**p*=0.0008, RFP vs *tai^RNAi^* \**p*=0.0002 (Unpaired student t-test with Welch’s correction).

Examining Dlp dynamics further, we find that RNAi depleting the extracellular protein Pentagone, which promotes Dlp/Dally glypican internalization and turnover[34], produces little change in Dlp levels among Tai-depleted cells (**Fig. S3A-C**, see bracketed regions). RNA-seq analysis of Tai-expressing wing discs detects elevated *dlp* mRNA, indicating that Tai may control *dlp* mRNA abundance (**Fig. S4)**. Consistent with this hypothesis, EcR CUT&RUN analysis [35] in 3^rd^ instar larval wing discs detects EcR binding peaks on the *broad* and *dlp* loci; *dlp* is flanked by two EcR peaks with another located in a *dlp* intron (**Fig. S4**). Orthogonal slices that traverse the Wg stripe of *en>tai* discs from control (anterior) to Tai overexpressing (posterior) regions confirm that Tai overexpression lowers Wg protein levels (**Fig. S5**). Notably, expression of a *dlp* transgene causes Wg accumulation on the DV flanks of Tai-depleted wing discs (**Fig. 7E-E’**), indicating that *tai* RNAi cells are similar to *wt* cells and retain Wg when provided extra Dlp[31]. However, expression of *dlp* together with *tai* (*en>tai,dlp*) has a reciprocal effect of enhancing Wg loss protein at the DV margin (**Fig 7F-F’**) and Wg build-up on the dorsal hinge (asterisk in **7G’**). Given that Tai expression with the same driver (*en>tai*) elevates Wg transcriptional output across the pouch (see **Figs. 4-5**), this loss of Wg may be due to enhanced trafficking and/or uptake. These data suggest that Tai status determines the effect of excess Dlp on Wg levels: Dlp depletes Wg levels across the pouch cells when excess Tai is present but retains Wg in Tai-depleted cells. A reciprocal test of a Dlp role in Tai overexpressing discs indicates that two different *dlp* RNAi lines do not rescue gaps in the Wg stripe that appear with Tai-overexpressing (arrows, **Fig. S6A-D**). These data indicate that Tai status determines the effect of excess Dlp on Wg, but that excess Tai may engage a different mechanism to modulate Wg protein in DV margin cells.

In the course of these analyses, we noted that wild type Tai or a version of the protein lacking PPxY motifs required to bind Yki [15] (Tai^PPxA^) drive accumulation of Dlp protein across the pouch (*enGal4*) (**Fig. S7A-B**) including in the DV midline region where Dlp is normally excluded (e.g., see arrows in **Fig. 7A**). Under normal circumstances, Dally is expressed in this region (e.g., see **Fig. S2**) and promotes short range Wg signaling from source cells[36]. Consistent with this local dependence on Dally, Tai-depletion only in DV midline cells (*bbg-Gal4*) has only a mild effect on distribution of Wg protein; however, providing exogenous Tai to these cells broadens the Wg-positive domain (**Fig. 8A-C**). Given that Tai does not elevate *wg* transcription (see **Fig. 5C**), this finding indicates that overexpressing Tai in DV midline cells is sufficient to enhance local spread of Wg protein and increase abundance of Wg-positive puncta (in **Fig. 8B’**) which have been shown to correspond to secretory vesicles[37]. Thus, expression of Tai in source cells enhances Wg secretion, while expression in the DV flanks can cooperate with excess Dlp to deplete Wg across the P-domain.

**Figure 8:**
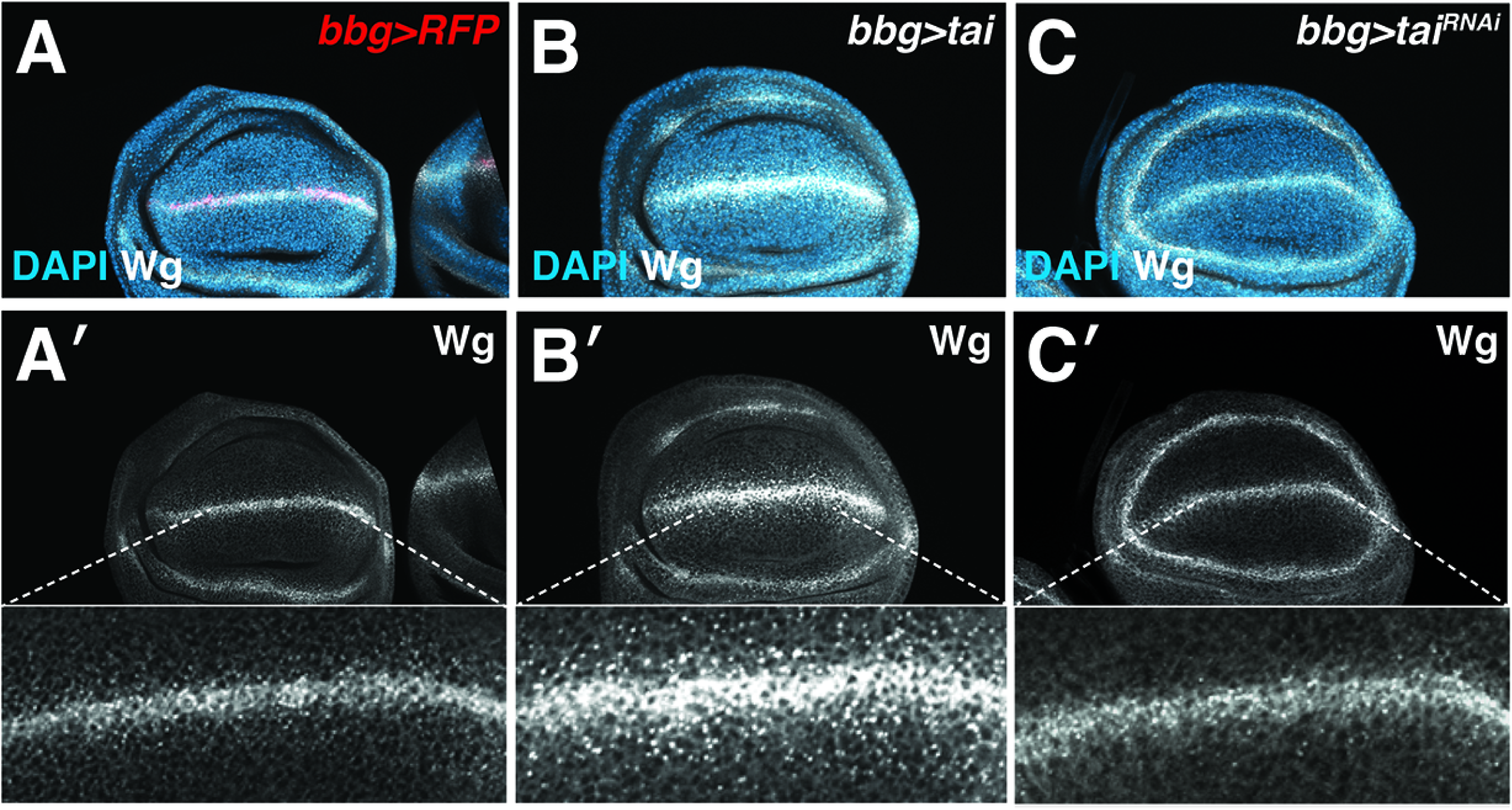
Tai expression at the D/V boundary enables excessive Wg spread. (**A-C**) Anti-Wg signal (greyscale) in **(A-A’)** *control* (*bbg>RFP*), (**B-B’**) Tai-overexpressing (*bbg>tai*), or Tai-depleted (**C-C’**) discs co-stained for nuclei (DAPI; blue). Dotted lines denote magnified views of Wg in **A’-C’**.

### Dlp protein is altered in *tai^low^* clones

Relative to the lethal *tai* RNAi transgene (e.g., with *actin5c-Gal4*[38]), the viable *tai^low^* allele may be predicted to yield correspondingly weaker phenotypes. To improve recovery of comparatively rare *tai^low^* pouch clones, *hsFlp* was applied in the *H99/+* background (see **Fig 3**). In this configuration, *tai^low^*pouch clones contain elevated levels of Dlp relative to adjacent control cells (arrows, **Fig. 9A-A”**). This result suggests that *tai^low^* could perturb Dlp intracellular traffic, either during secretion or internalization. To test this model, Dlp was reanalyzed in *tai^low^* clones under conditions that do not permeabilize membranes; this approach detects only basal levels of Dlp on the surface of control and *tai^low^* disc cells (**Fig. 9B-B”**). Indeed, closer inspection of *tai^low^* hypomorphic clones in tangential slices indicates that Dlp is mislocalized to the cytoplasm (**Fig. 9C-D**). Given that removing Pent, which stimulates Dlp internalization[34], does not rescue Dlp loss in Tai-IR cells, Tai may promote both *dlp* mRNA expression and Dlp protein secretion, and that the consequent decline in ability to compete for Wg with normal neighbors deprives *tai^low^* cells of a signal necessary for growth and survival.

**Figure 9:**
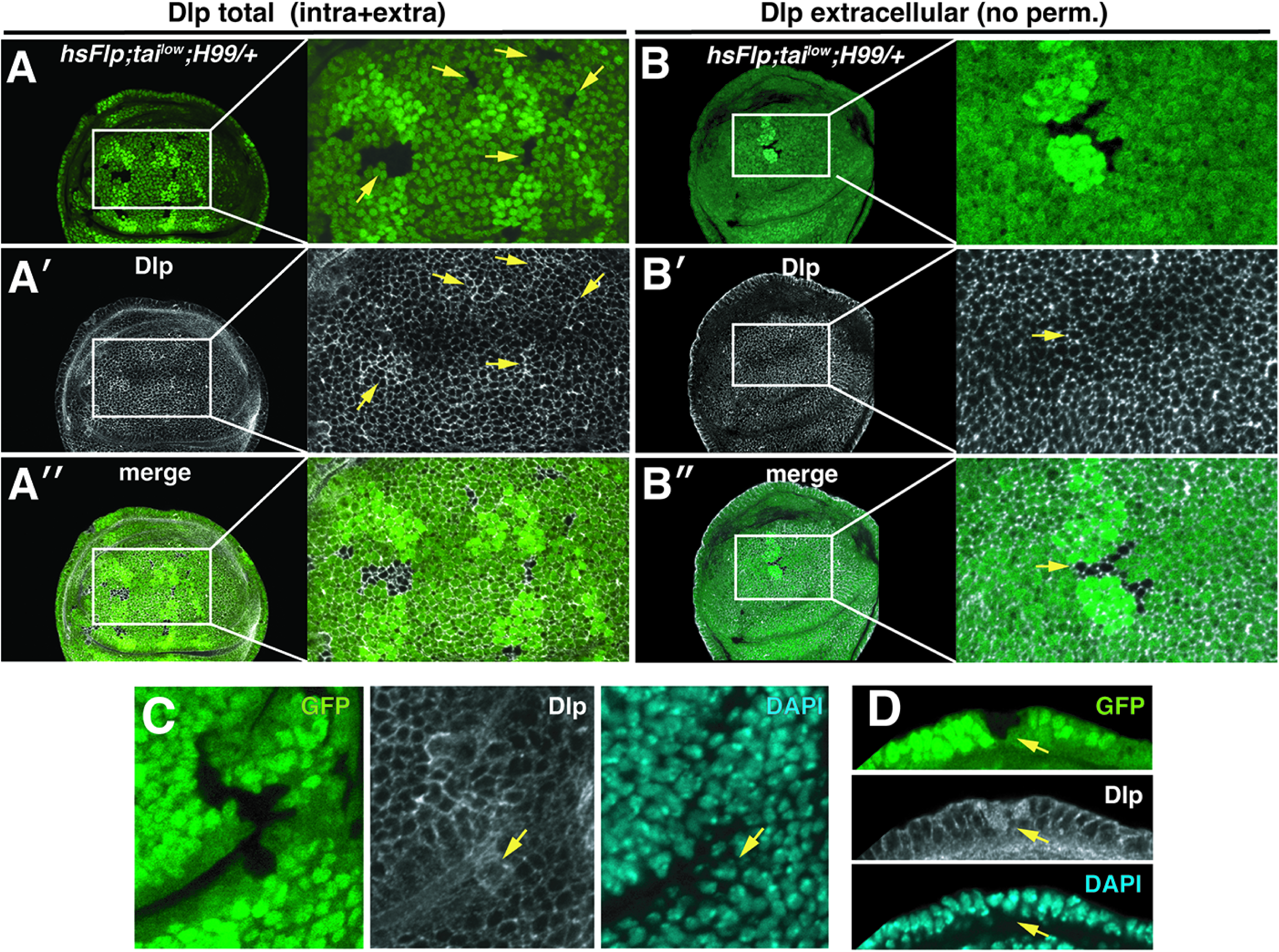
The *tai^low^*hypomorph causes Dlp intracellular accumulation in pouch cells. (**A-D**) *hsFlp* generated *tai^low^* clones (GFP negative) in a *H99/+* background co-stained for Dlp (greyscale) and nuclei (DAPI; cyan). Discs in panels **A-A”** and **C** were permeabilized to detect total Dlp (intracellular and extracellular), while discs in panel **B-B”** were processed without permeabilization and only detect extracellular Dlp. Arrows in **C** and **D** denote *tai^low^* cells in side-view with intracellular Dlp.

## DISCUSSION

Here we show that cells with a reduced the dose of the Tai coactivator, which functions in the ecdysone and Hippo pathways[15,39], are killed by normal neighbors through a mechanism involving competition for the Wingless (Wg/Wnt) ligand. Elevated Wg signaling significantly rescues elimination of Tai^low^ cells in multiple organs, suggesting that Tai may normally promote Wg activity. Examining distribution of Wg components reveals that Tai promotes extracellular spread of the Wg ligand from source cells across the wing disc, thus ensuring patterned expression of multiple Wg-regulated target genes. Tai controls Wg spread indirectly through the extracellular glypican Dally-like protein (Dlp), which binds Wg and promotes its extracellular diffusion and capture by receptors. Data indicate that Tai likely controls Dlp at two levels: transcription of *dlp* mRNA and Dlp intracellular trafficking. Overall, these data indicate that the Tai acts through Dlp to enable Wg transport and signaling, and that cell competition in the Tai^low^ model arises due to inequity in the ability of epithelial cells to sequester limiting amounts of the Wg growth factor.

One key conclusion from these data is the mechanism through which Tai^high^ cells kill normal neighbors (e.g., through Spz-Toll)[6] is distinct from the mechanism that eliminates Tai^low^ cells. The former mechanism is produced by triggering an innate immune pathway and may require priming by chronic microbial infection[18], while the latter derives from a developmental role in Dlp-mediated Wg diffusion and turnover (this study). These mechanisms may both occur in vivo, although in different contexts, with Tai^high^ overexpression mimicking oncogene activation in cancer, and Tai^low^ representing cells in developing tissues that receive weaker proliferative or survival signals than neighbors. In the developing wing disc, gradual scaling of the Wg gradient as the tissue grows ensures that adjacent cells experience only incremental changes in concentration of pro-growth morphogens[40]. Artificially steepening of a gradient, as occurs in *pent* mutant wing discs[21], or creating patches of cells deficient in morphogen capture (e.g., with *tai^low^* or *tai^RNAi^*) or signaling (e.g., as in Wg pathway mutants[17]) within an otherwise intact gradient each create sharp differences in morphogen activity. This model is supported by the finding that artificial boundaries between glypican expressing and non-expressing cells is sufficient to induce cell competition among *Drosophila* follicle stem cells[41]. Intriguingly, *dlp* follicle clones are ‘winners’ while *dally* clones are ‘losers’, which implies different roles for Wg in survival of pouch cells vs. follicle stem cells.

The finding that *tai* alleles perturb Dlp expression, retard the Wg gradient, and interact with *Apc* alleles in eye and wing epithelia seems to suggest that Tai gain and loss phenotypes in these tissues may be driven in part by changes in Wg signaling. It remains to be determined whether Tai acts broadly across multiple tissues and developmental stages to modulate Wg, or whether this role is limited to specific cell types. As Tai is a transcriptional coactivator it is likely to mediate its Wg regulatory effects through other factors that directly contact DNA. The identity of this factor(s) is unclear. One candidate is the ecdysone receptor EcR, which brings Tai into a transcriptional regulatory complex whose components are conserved across metazoans (i.e., ERα-NCOA3-p300)[42]. The role of EcR in innate immune transcription[43] is consistent with our finding that EcR is involved in intertissue invasion led by Tai^high^ cells[6]. However, the role of an EcR-Tai complex in death of Tai^low^ cells has not been examined. Ecdysone can promote (or ‘gate’) activity of both major morphogen gradients in the wing disc[44], and the presence of EcR CUT&RUN peaks in the *dlp* genomic region (see **Fig. S4**) seems to imply EcR may have some role in Dlp expression. EcR can also repress *wg* transcription in areas of the pouch[45], much as we have found that Tai represses *wg* transcription on either side the DV margin (see **Fig. 5B**). However further work is required to determine whether either of these roles requires an EcR-Tai complex. Mild rescue of *tai^low^*elimination by a *wts* allele indicates that slight elevation of Yki activity can promote *tai^low^* survival. However, the ability of the non-Yki binding form of Tai, Tai^PPxA^, to efficiently induce Dlp accumulation in pouch cells is not consistent with a requirement for Tai to bind Yki during the competition process. Alternatively, Tai may modulate Wg signaling independent of interactions with EcR and Yki, perhaps through other transcription factors. Determining Tai-binding partners involved in regulation of the Wg pathway and establishing Tai transcriptional targets whose products modulate secretion and/or endocytosis of glypicans could yield broad insight into mechanisms that calibrate morphogen diffusion in development and disease.

### Lead contact and resource availability

Requests for resources and reagents should be directed to and will be fulfilled by Lead Contact.

## Acknowledgements

We thank the Emory Integrated Cellular Imaging Core for their expertise and assistance. We also thank E. J. M. Storkebaum, the Developmental Studies Hybridoma Band (DSHB), Bloomington Drosophila Stock Center (BDSC), the Harvard Transgenic RNAi Project (TRiP), and the Vienna Drosophila Stock Center for antibodies and stocks. We also thank members of Moberg laboratory for their helpful discussion. This work was funded by NIH T32 grants GM008367 (CSK) and T32 GM149422 (VCP), F31 Predoctoral Fellowship CA254207 (CKS), and GM121967 (KHM)

## Author Contributions

Conceived and designed the experiments: CKS, KHM. Performed the experiments: CKS. Analyzed the data: CKS, VCP, KHM. Authored the paper: CKS. Edited the manuscript: CKS, KHM. Funding and Acquisition: CKS, KHM.

## Materials and Methods

### Drosophila culture

Fly stocks were maintained under standard culture conditions at 25°C, 12 hr light:dark cycles in humidity-controlled incubators. Experiments used a mix of female and male animals. All crosses were maintained on standard molasses food supplemental with yeast. Unless specified, flies were mated and allowed to lay eggs for a period of 24-48 hours, and then dissected at wandering third instar. Lines used are referred to in **Key Resources Table** (BDSC stock numbers indicated).

### Strategy for randomization and/or stratification

Animals of both sexes were used for experiments. A minimum of 10 total imaginal wing discs over three biological replicates were used in all experiments. Figures presented are representative images of each genotype.

### Adult eye screen

Adult females were aged for three days to allow condensation of pigment granules then frozen at -20°C for >5 minutes before imaging with a Nikon SMZ800N microscope at 25X magnification (2.5X objective x 10X eyepiece) using a Leica MC170HD camera and a Nii-LED High intensity LED illuminator. Flies were submerged in 70% ethanol to eliminate glare. Multiple focal planes were imaged and merged in Photoshop to generate a final image.

### Cell Competition Assays

Mated flies were allowed to lay for 24 hrs before flipping into a new vial. 48-50 hrs after egg lay (AEL), FLP recombinase was induced by heat-shocking of the larvae at 37°C for about one hour, except for in the generation of “Flp-out” clones were larvae were heat-shocked for 1.5 hrs. Third instar larvae were then dissected at 96 hrs AEL. Both male and female larvae were dissected. Clonal areas were measured using ImageJ, and only clones in the wing pouch were considered for analysis.

### Immunofluorescence and microscopy

Immunofluorescence and microscopy were performed using standard methods. Briefly, wing discs were dissected in 1X phosphate-buffered saline (PBS), fixed for 20 minutes in 4% paraformaldehyde (PFA) at room temperature (RT), rinsed 5X in 1X PBS, and then permeabilized with 1X PBS and 0.3% Triton X-100 (0.3% PBST). For extracellular stain without permeabilization, the protocol was performed as in [46]. Discs were then blocked with 10% normal goat serum (NGS) and then incubated in diluted primary antibody containing 10% NGS and 1X PBS and 0.1% Triton X-100 (0.1% PBST). Samples incubated in primary antibody solution at 4°C overnight, washed 5X with 0.1% PBST, incubated with DAPI (1:500) for 5 minutes, washed 3X with 0.1% PBST, and then incubated for 1 hr at RT with secondary antibody solution diluted with 10% NGS and 0.1% PBST. Discs were then washed 5X times with 0.1% PBST and incubated in n-propyl gallate (NPG, 4% w/v in glycerol) overnight. Discs were mounted in NPG on glass coverslips and then imaged with a Nikon A1R confocal system using 20X and 40X objectives. Images were processed with Fiji, Photoshop, and Illustrator software. ImageJ was used for clonal area measurements. Refer to **Key Resources Table** for full list of antibodies, dilutions, and other reagents used.

### Fluorescence intensity analysis

Fluorescence intensities were measured across equal regions of interest (ROIs) using Image J/Fiji. Briefly, the ROI was positioned such that it was: 1) centered over the AP boundary as defined by the *en>GFP* control channel, 2) below the DV line (approximately halfway between the DV boundary and the bottom of the pouch), and 3) extended approximately halfway into each compartment (anterior and posterior). Raw fluorescence intensity values were standardized such that the mean anterior fluorescence was equal to 1, and posterior fluorescence was relative to anterior. Analyses and fluorescence intensity plots were performed and created in R (R version 4.1.1 (2021-08-20)). Posterior: anterior mean fluorescence intensity ratios were plotted, and statistics performed using GraphPad Prism. Unpaired t-tests with Welch’s corrections were performed.

### Quantitation and Statistical Analyses

Unpaired Student’s t-tests or one-way ANOVAs (GraphPad Prism) were used to analyze significance between datasets. Significance was considered to be *p*<0.05. Normal distributions were assumed.

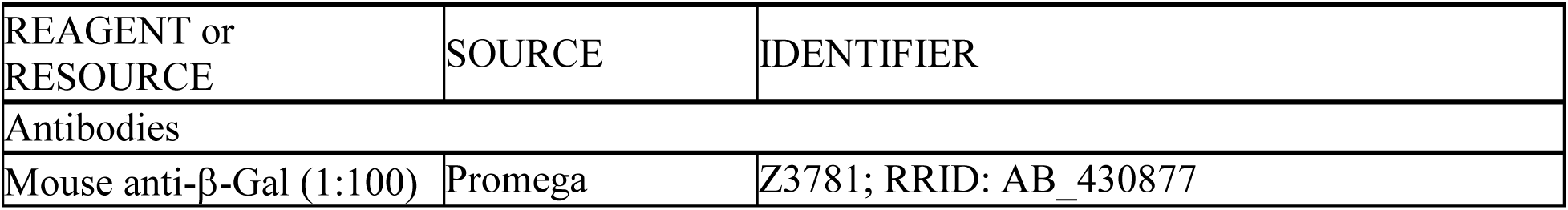
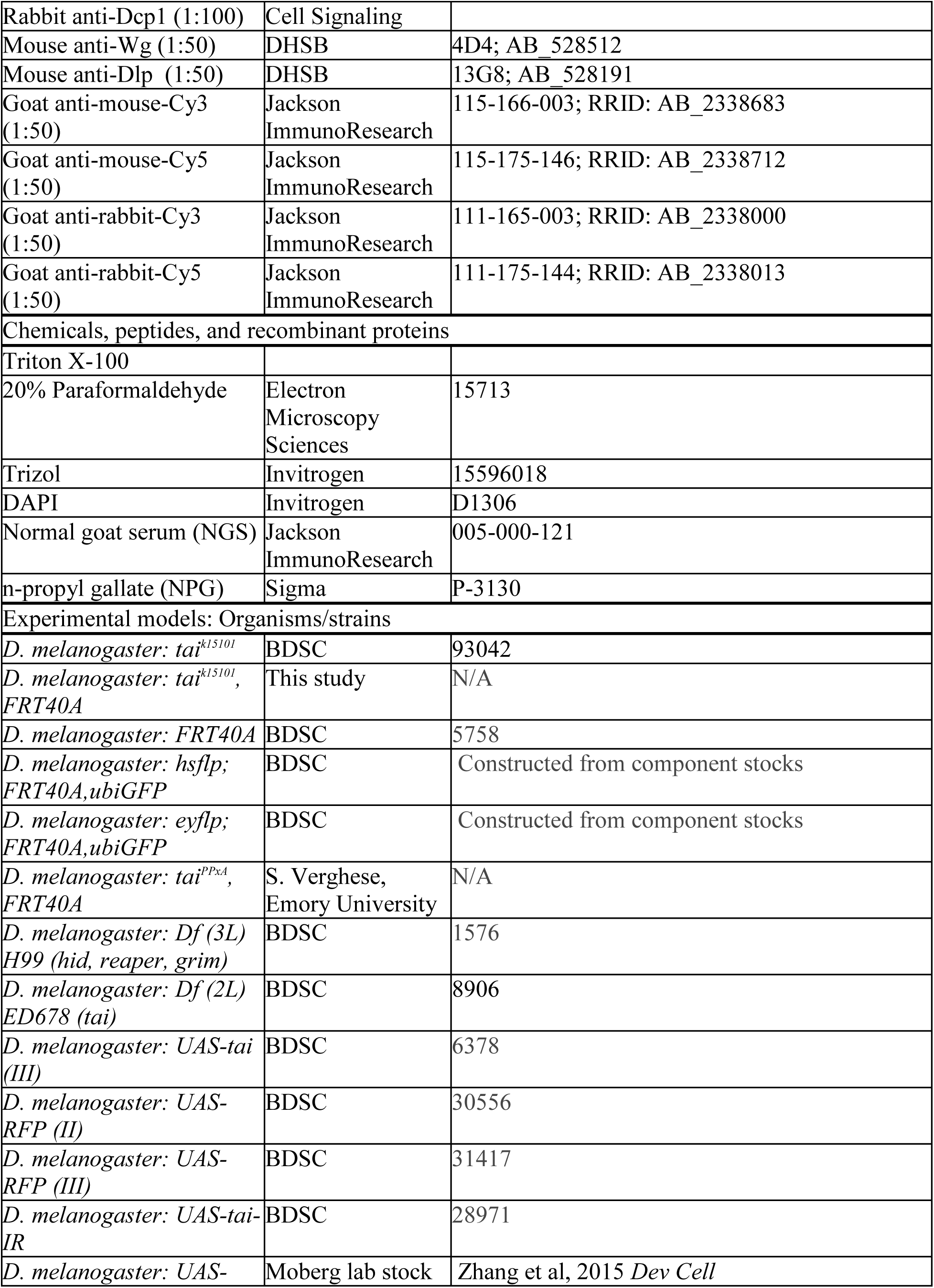
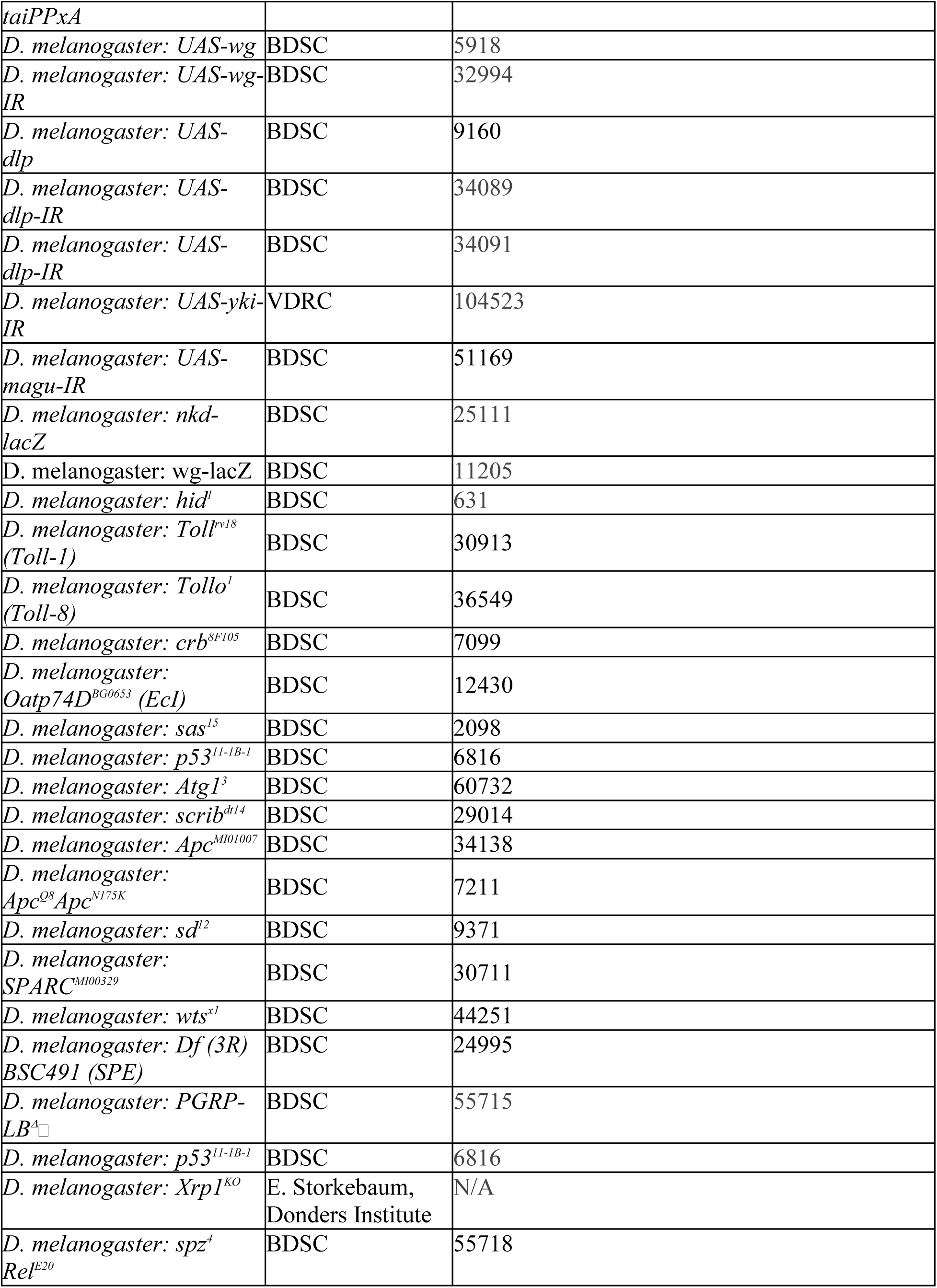
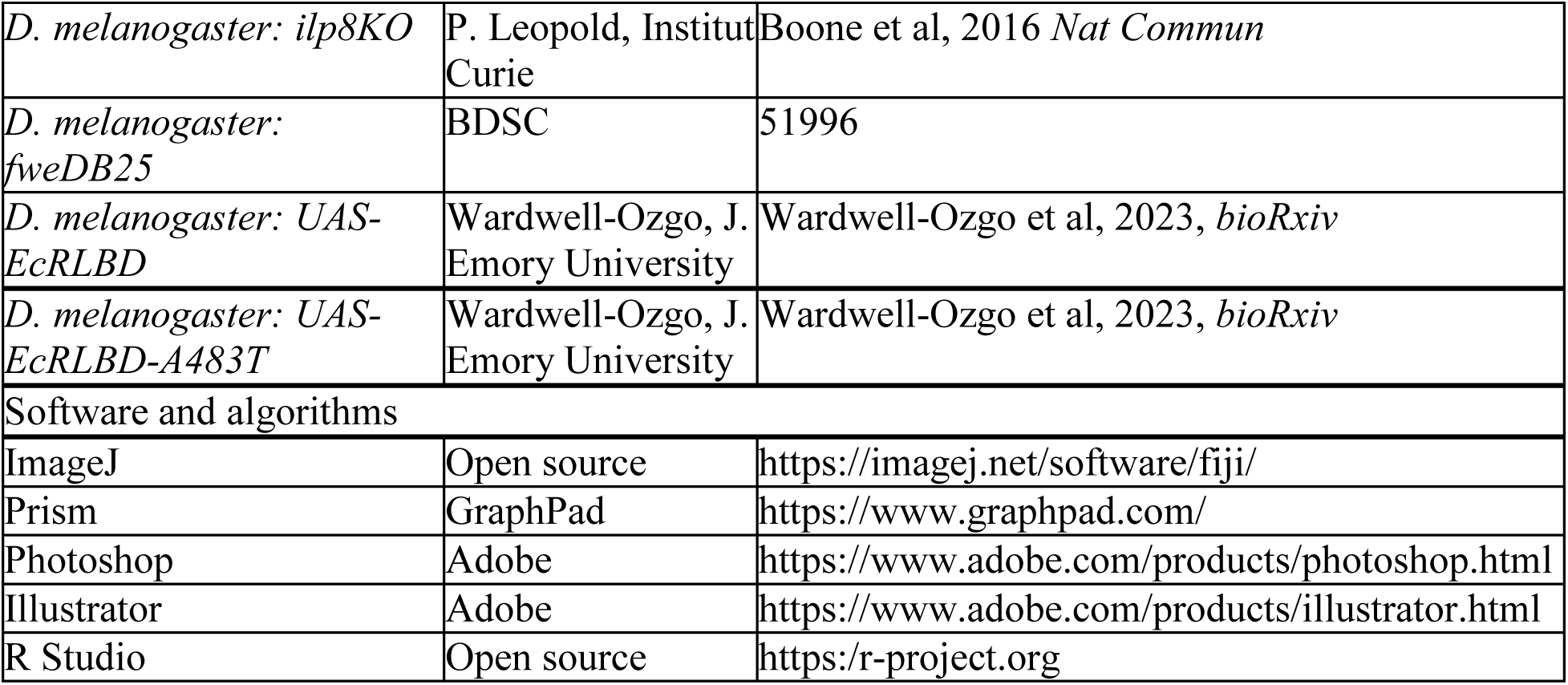
KEY RESOURCES TABLE

## Supplemental Figure Legends

**Figure S1. *H99* and *Apc* alleles allow *tai^low^*cells to survive 76hrs after clonal induction.** Wing discs bearing 76hr old (post-hs) pouch clones of **(A)** *FRT40A* cells or (**B-G**) *tai^low^* cells in the indicated genetic backgrounds. Clones are GFP-negative and twinspots are bright green. (**H**) Clone:twinspot area ratios (pixels) of the same genotypes as in **A-G.** Note significant rescue of *tai^low^* survival by *H99* and *Apc* heterozygosity. \*\*\*\**p* <0.001 ***p = 0.0007, ns = not significant (One-way ANOVA with Dunnett post hoc test).

**Figure S2. *dally* does not respond to Tai depletion but can be induced by excess Tai**. (**A-C**) Expression of the *dally-lacZ* reporter detected by anti-ßgal stain (greyscale) in (**A-A”**) *control* (*en>GFP,RFP*), (**B-B”**) Tai overexpressing (*en>tai,GFP*), or (**C-C”**) Tai-depleted (*en>tai^RNAi^,GFP*) discs. Arrows in **A’** and **C’** indicate shrunken pattern of *dally-lacZ* expression in a Tai-depleted P-domain.

**Figure S3. RNAi of Pent, which promotes Dlp internalization, does not fully restore Dlp in Tai-depleted disc regions.** Anti-Dlp signal in (**A-A”**) *control* (*en>GFP*), (**B-B”**) Tai-depleted (*en>tai^low^*) discs, or discs (**C-C”**) co-depleted for Tai and Pent (*en>tai^RNAi^*+*pent^RNAi^*). Brackets denote P-domain with Dlp depletion with *tai* RNAi.

**Figure S4. Evidence of *dlp* transcription regulated by Tai. Top:** IGV traces of RNA-seq reads from *control* (green; *en>GFP*) and Tai-expressing (red; *en>tai*) larval wing discs laid over (**middle**) a genomic map of the *dlp* region. **Bottom**: EcR-association peaks in the *dlp* region ().

**Figure S5. Excess Tai downregulates Wg levels within source cells at the DV margin.** Wg signal (greyscale) in a Tai overexpressing disc (*en>tai,GFP*) imaged *en face* (**A-A’**) and (**B-B’**) tangential slices corresponding to **region 1** and **region 2 in** magnified view **(B-B’**). Panel **C** and associated **regions 1** and **1** show equivalent analysis in *control* (*en>GFP,RFP*) discs.

**Figure S6. RNAi depletion of Dlp is not sufficient to block the Tai-induced gaps in the DV stripe of Wg.** Wg signal (greyscale) in wing discs expressing either of two *dlp* RNAi constructs alone (#1 or #2 in **A-B**) or in combination with Tai overexpression (**C-D**). Arrows in **C’** and **D’** mark gaps in the Wg stripe.

**Figure S7. Tai^PPxA^ is able to induce Dlp expression in the midline region.** Dlp signal (greyscale) in (**A-A”**) *control* (*en>GFP,RFP*) or (**B-B”**) Tai^PPxA^ expressing (*en>tai^PPxA^,GFP*) discs. Yellow arrows in **A’** mark Dlp exclusion from cells in either side of the DV margin; arrow in **B’** points to excess Dlp filling this region.

**Supplemental Table 1: A genetic screen reveals dominant modifiers of *tai^low^* cell death.** Heterozygous candidate alleles selected from various pathways were placed in the background of a mosaic *eyFlp; Tai^low^* eye, and positive screen “hits” were determined based on increased recovery of Tai^low^ clones (orange). An allele of *hid*, as well as the *H99* deletion (*reaper, hid, grim*) enhanced recovery of Tai^low^ cells. Other suppressors included *wts^x1^*, a loss of function allele of the core Hippo pathway kinase and an allele of *Adenomatous polyposis coli* (*Apc*), a key inhibitor of the conserved Wg/Wnt pathway. Refer to Key Resources Table for information about alleles used.

