## Supplemental Files 1-7 for "The *Drosophila* EcR-Hippo component Taiman promotes epithelial cell fitness by control of the Dally-like glypican and Wg gradient"

*hsFlp;FRT40*

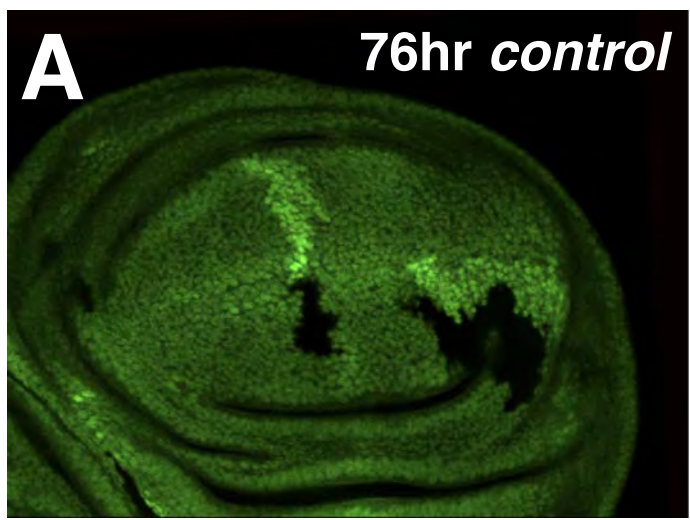

*hsFlp;tai<sup>low</sup>,FRT40A*

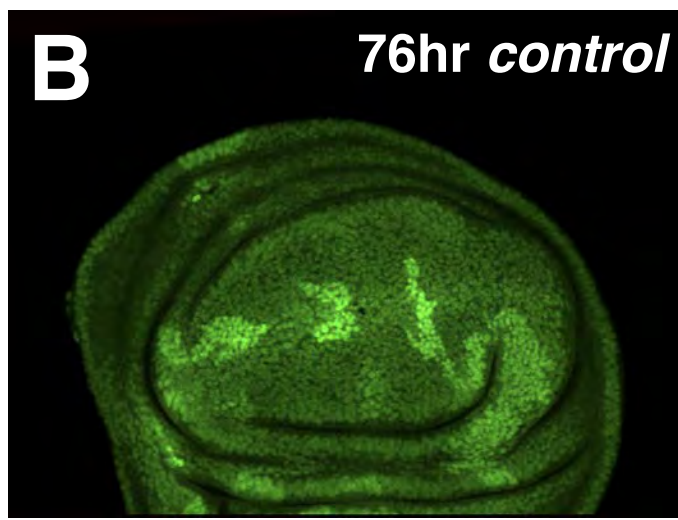

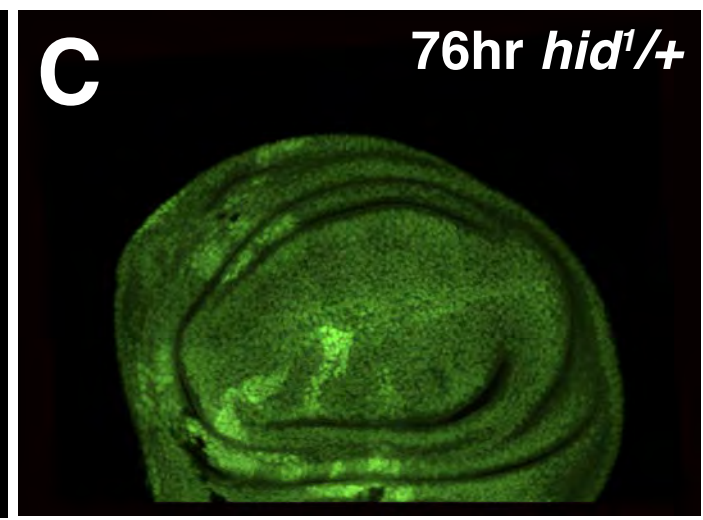

*hsFlp;tai<sup>low</sup>,FRT40A*

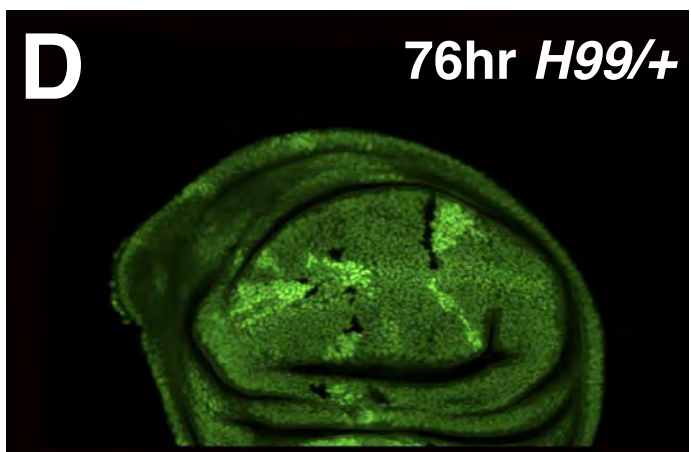

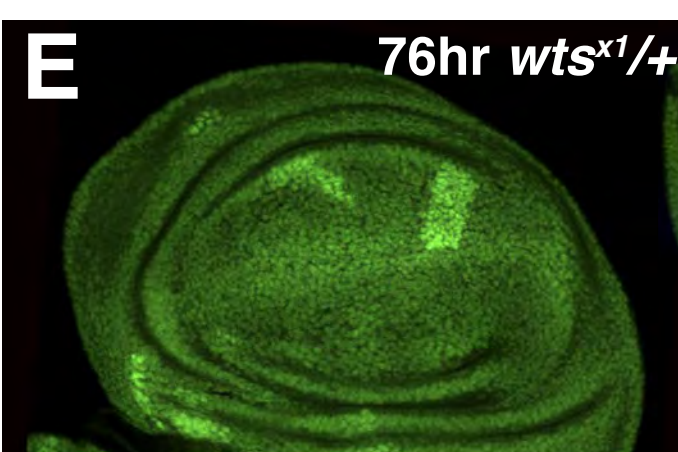

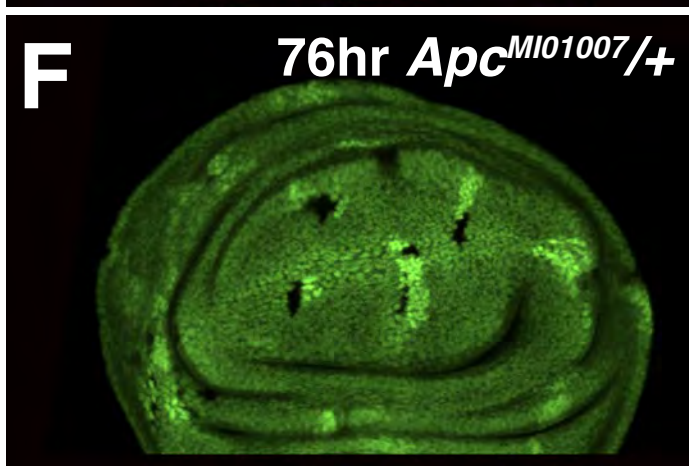

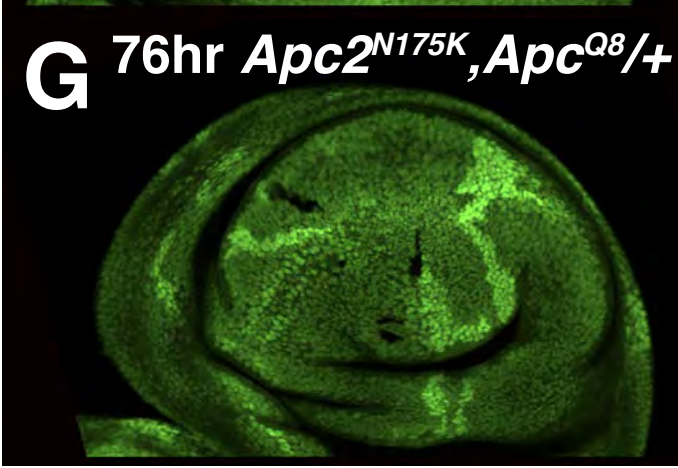

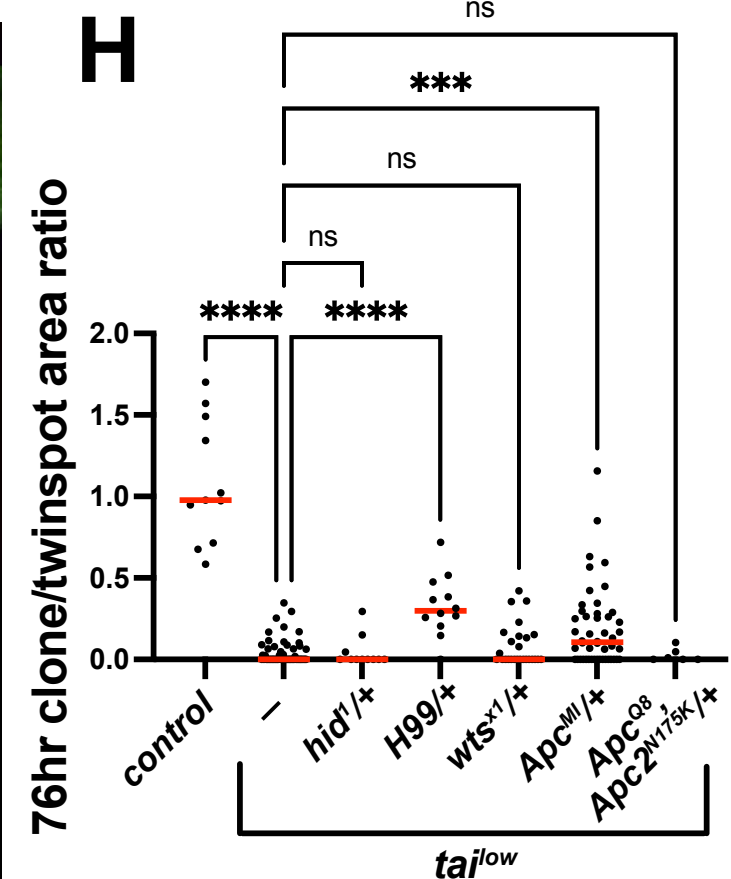

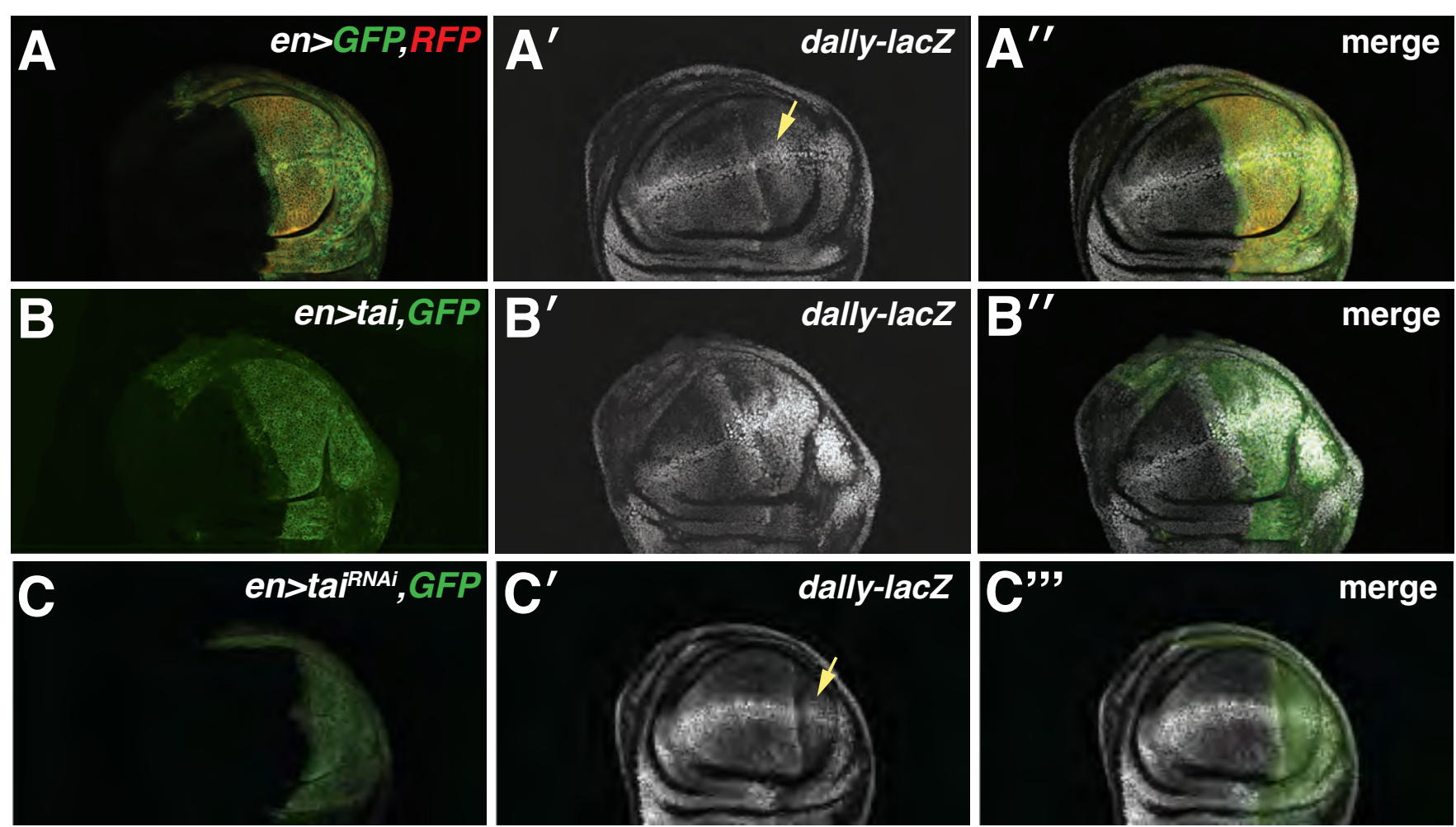

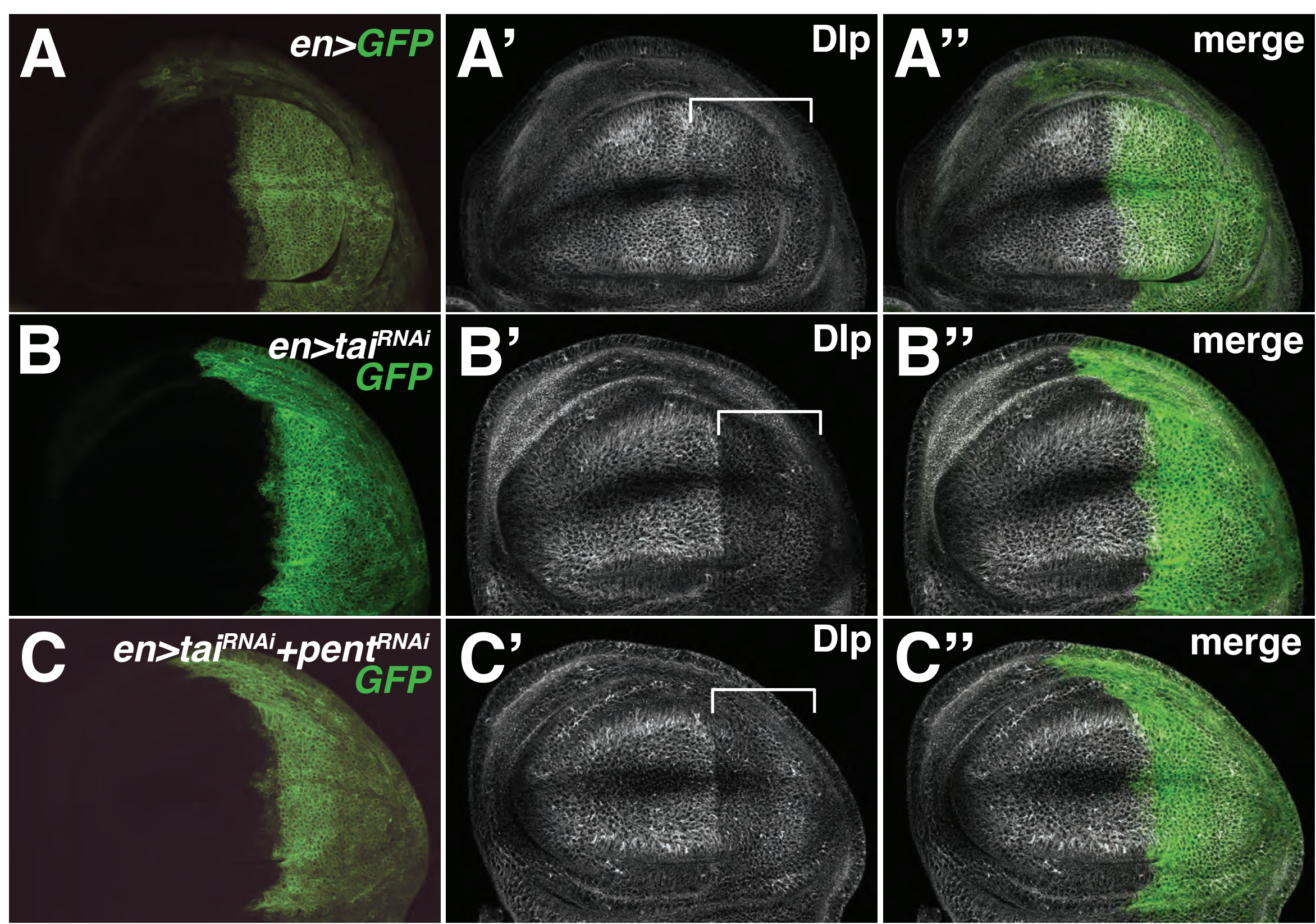

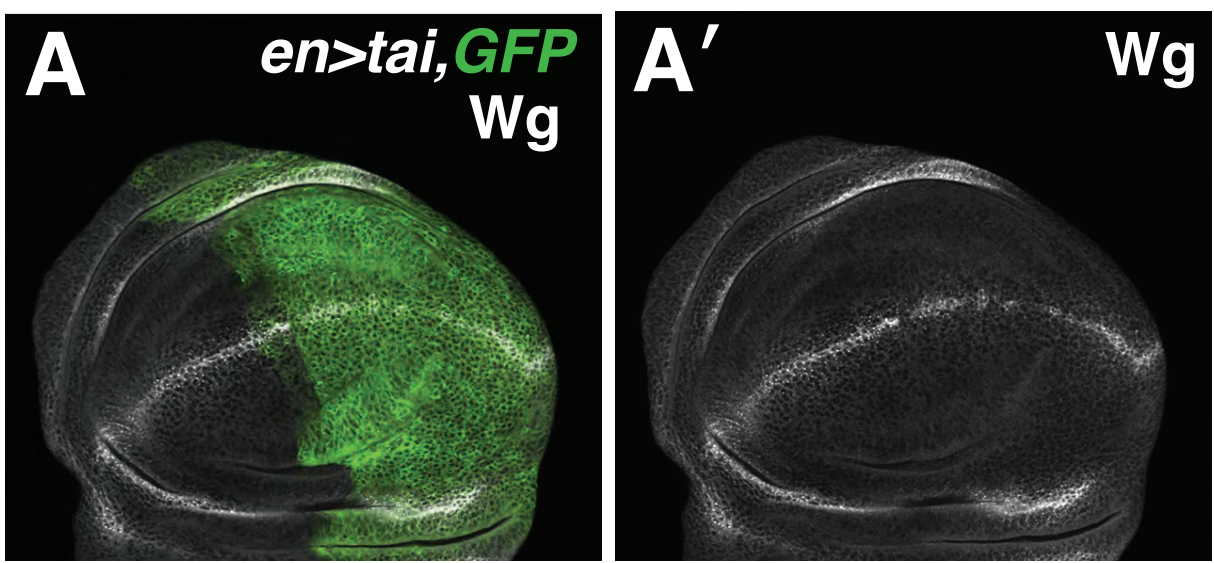

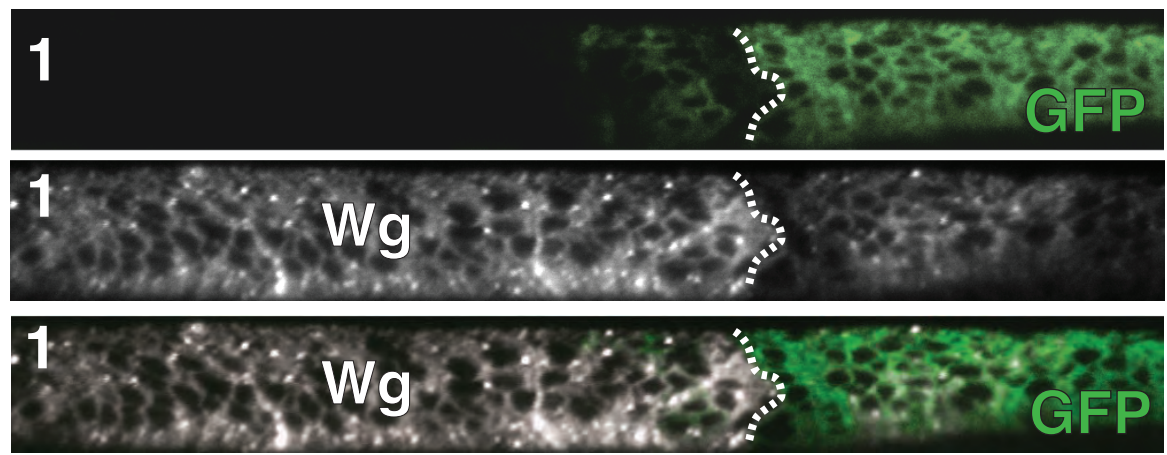

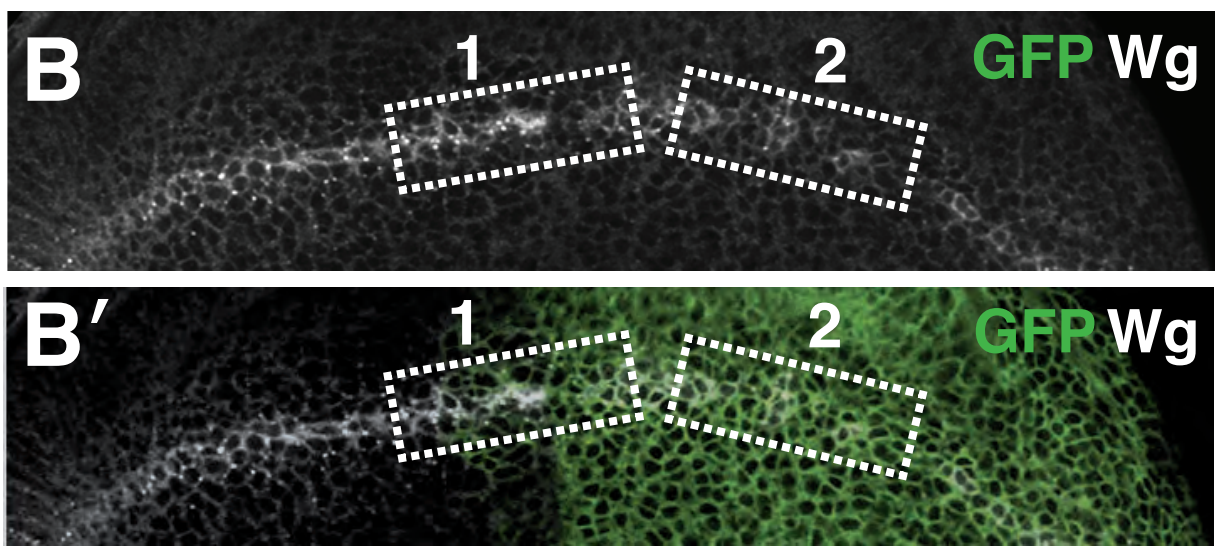

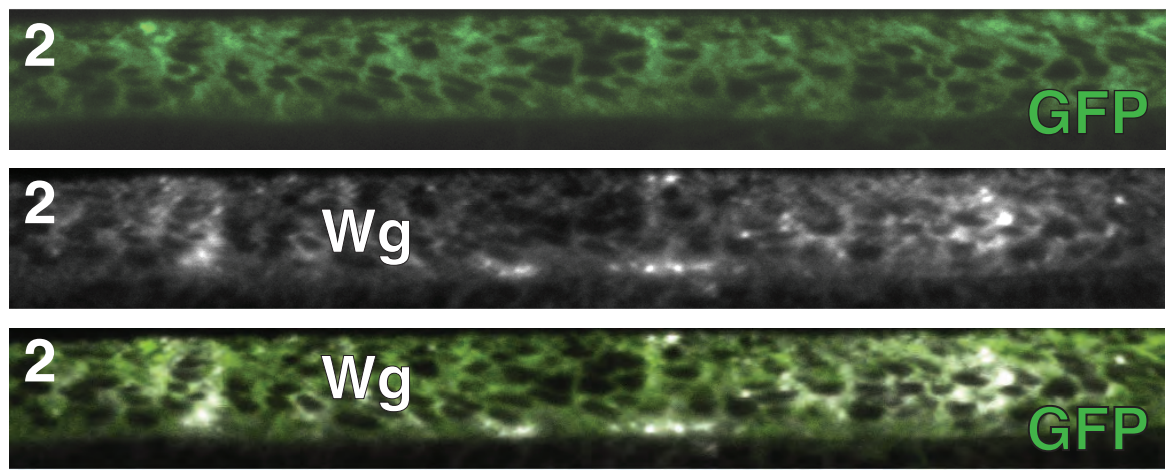

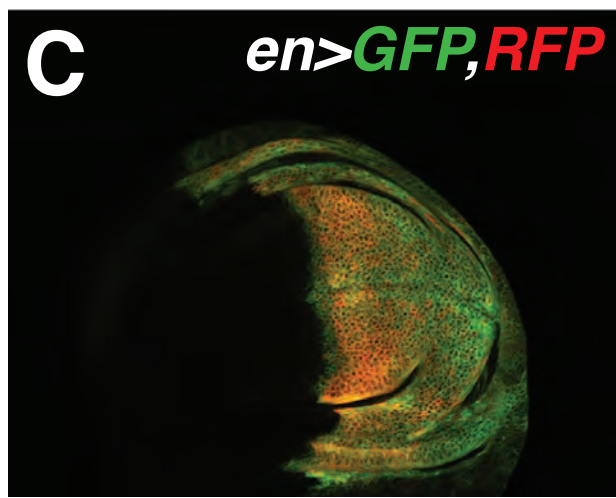

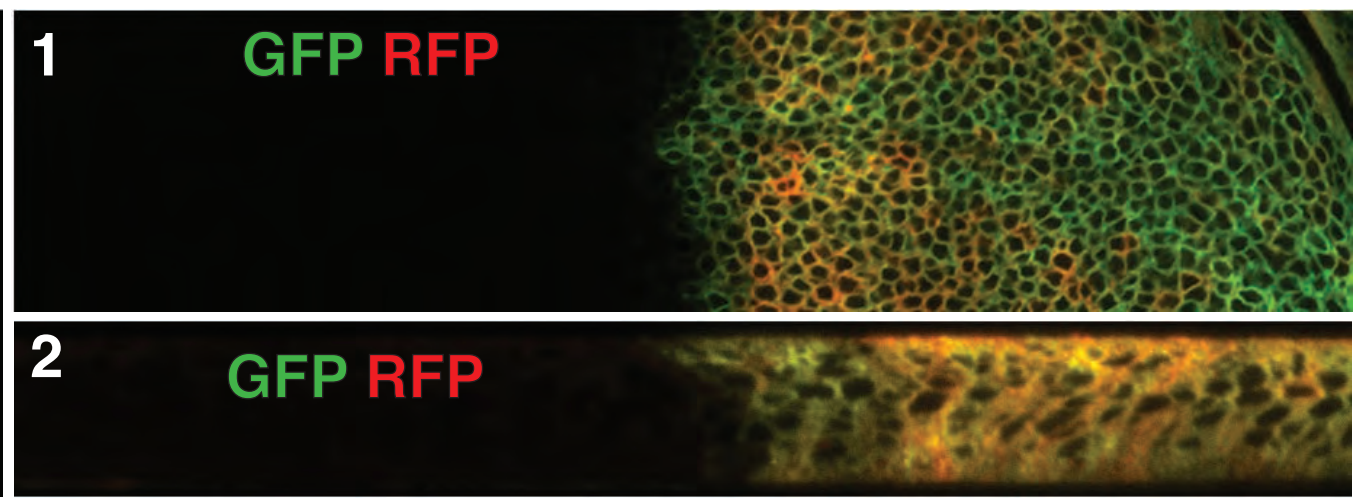

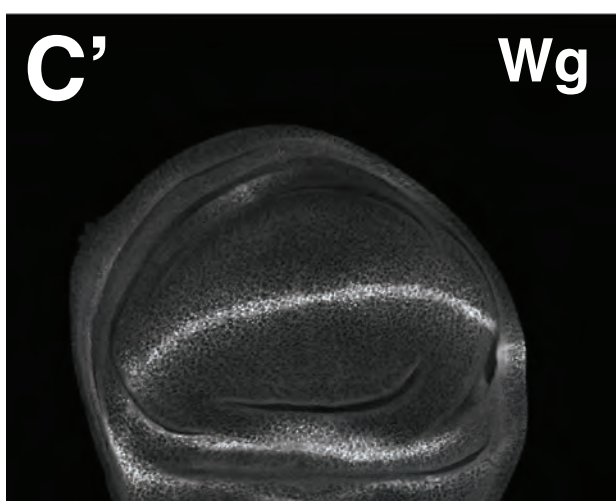

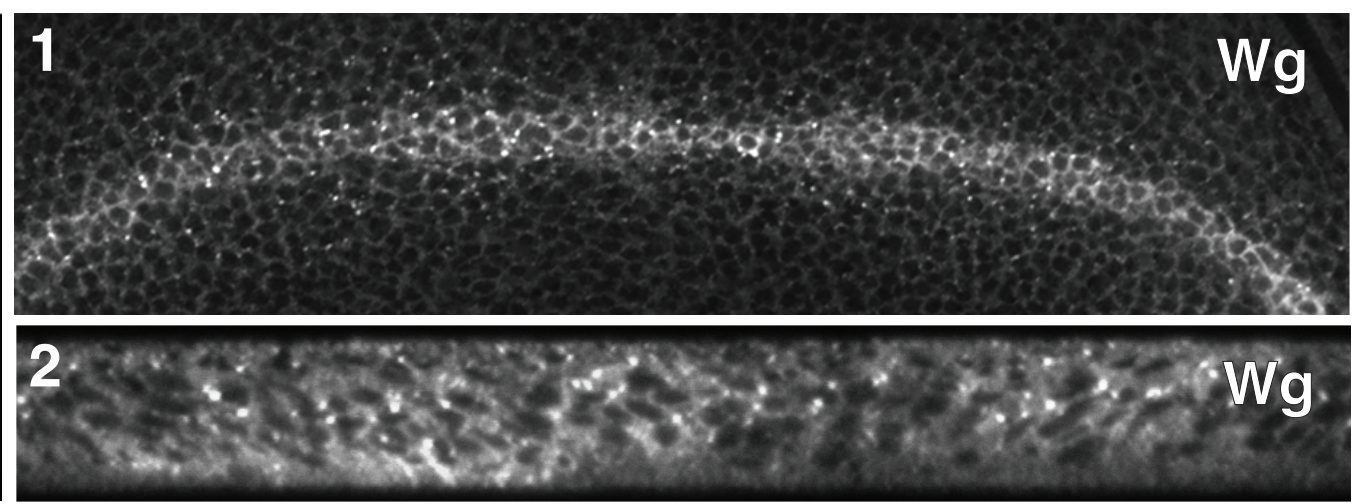

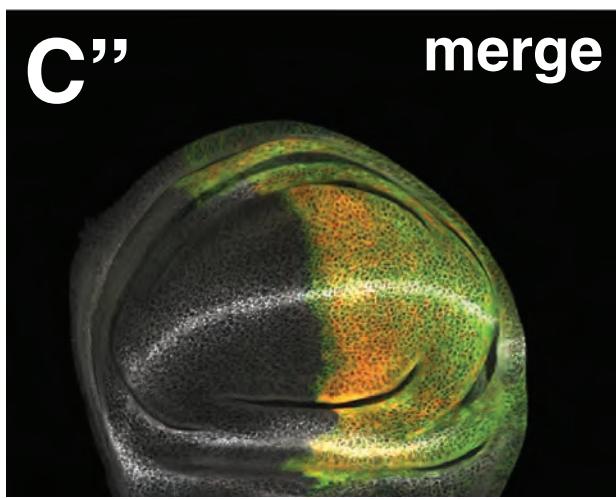

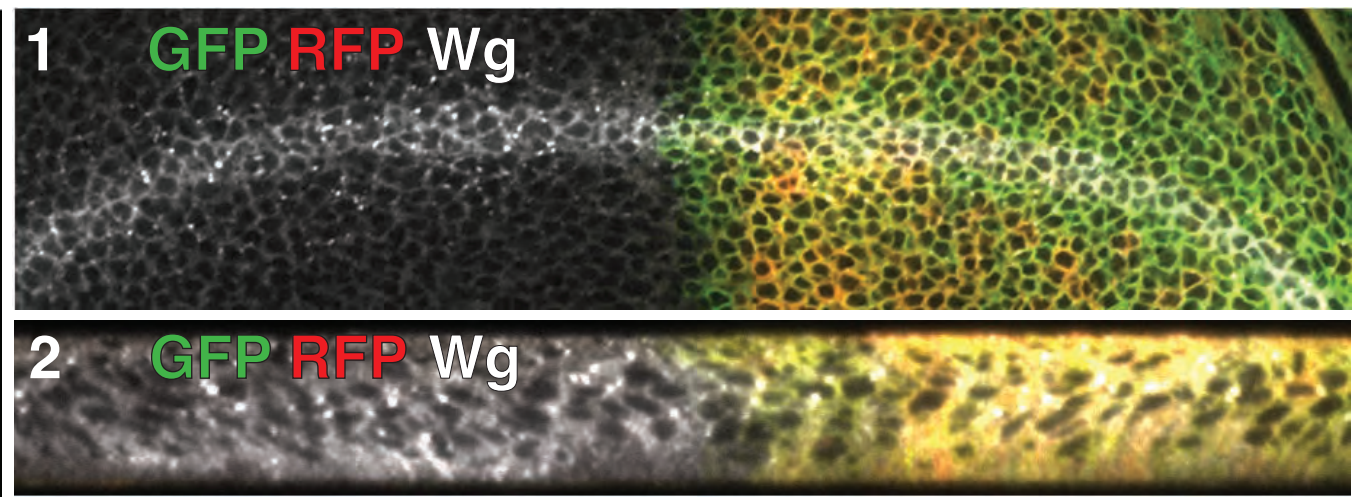

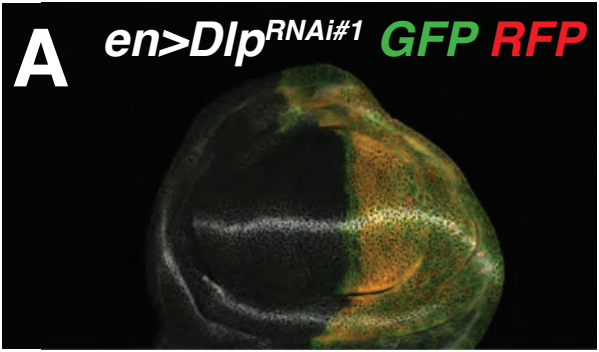

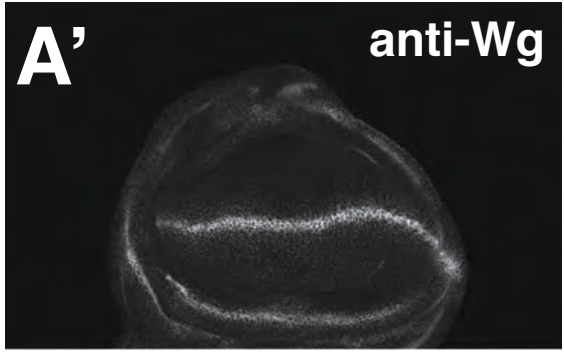

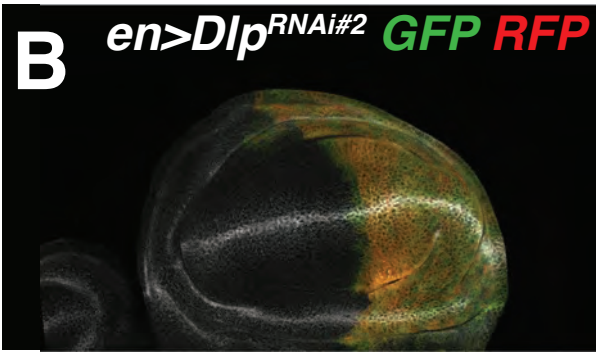

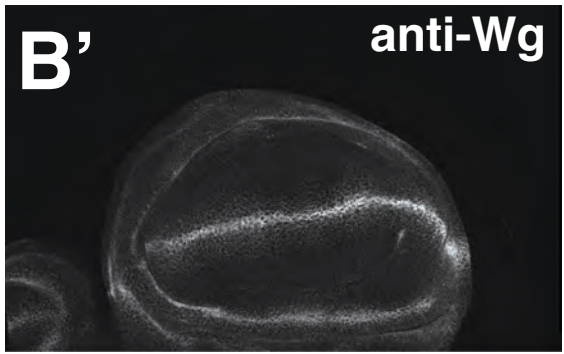

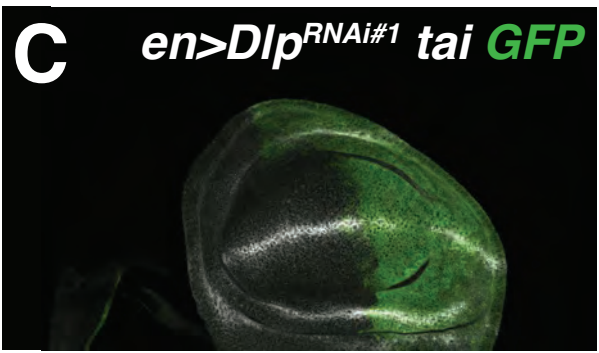

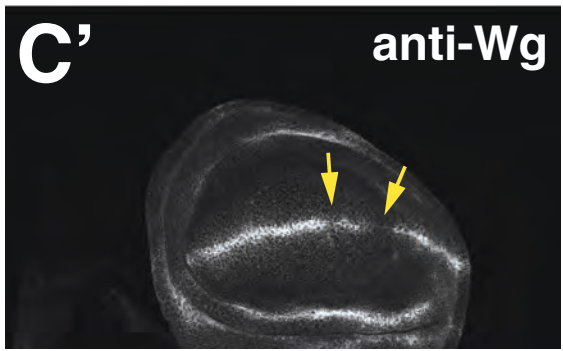

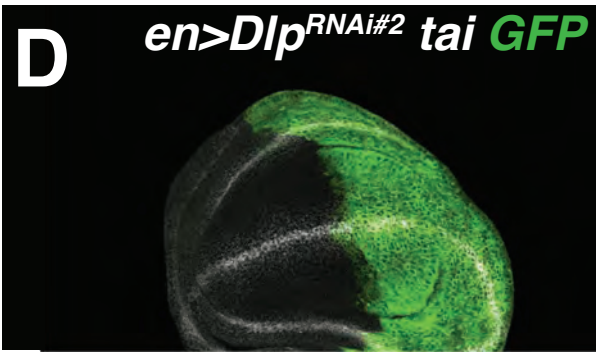

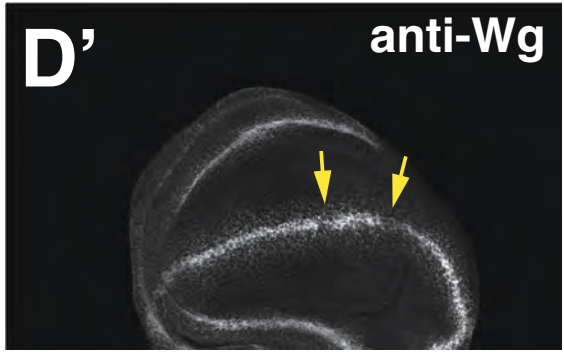

**A**

*en>GFP,RFP*

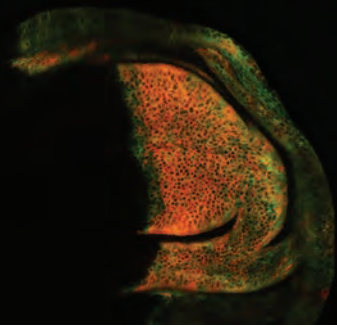

**A'**

Dlp

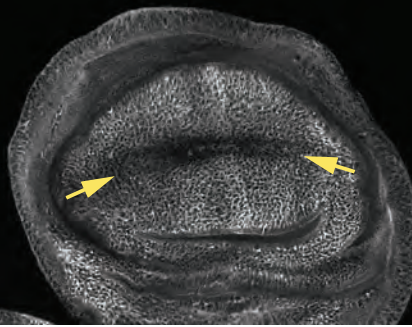

**A''**

merge

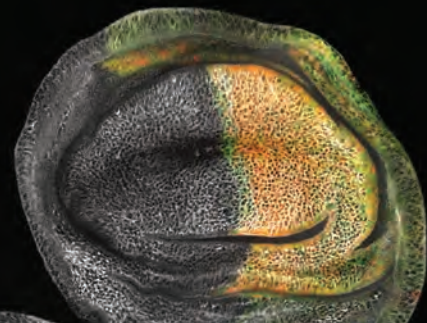

**B**

*en>tai<sup>PPxA</sup>,GFP*

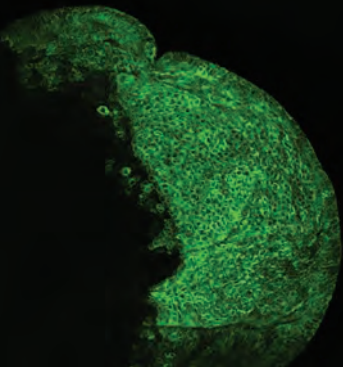

**B'**

Dlp

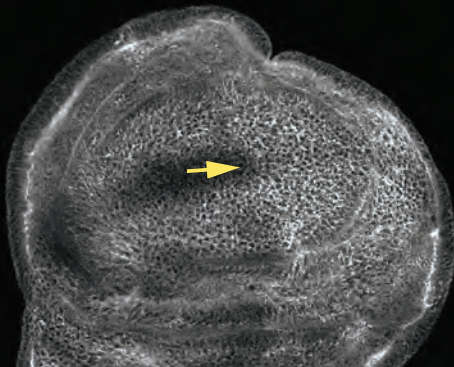

**B''**

merge

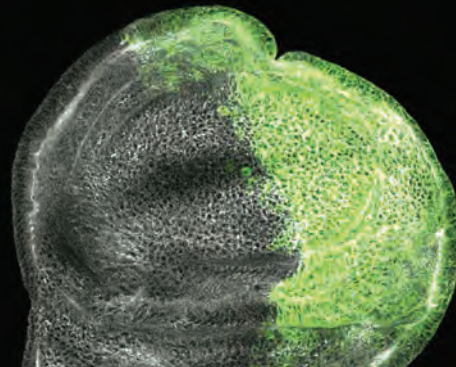
